# A modular system for programming multistep activation of endogenous genes in stem cells

**DOI:** 10.1101/2024.09.17.613466

**Authors:** Anupama K. Puppala, Andrew C. Nielsen, Maureen R. Regan, Georgina E. Mancinelli, Renee F. De Pooter, Stephen Arnovitz, Caspian Harding, Michaele McGregor, Nikolas G. Balanis, Ryan Clarke, Bradley J. Merrill

## Abstract

Although genomes encode instructions for mammalian cell differentiation with rich syntactic relationships, existing methods for genetically programming cells have modest capabilities for stepwise regulation of genes. Here, we developed a sequential genetic system that enables transcriptional activation of endogenous genes in a preprogrammed, stepwise manner. The system relies on the removal of an RNA polymerase III termination signal to induce both the transcriptional activation and the DNA endonuclease activities of a Cas9-VPR protein to effect stepwise progression through cascades of gene activation events. The efficiency of the cascading system enables a new dimension for cellular programming by allowing the manipulation of the sequential order of gene activation for directing the differentiation of human stem cells.

**One-Sentence Summary:** Development of a synthetic biology system for preprogrammed, stepwise activation of endogenous genes.

## Introduction

Cellular differentiation is a sequential process, in which progenitor cells progress through successive intermediate cell states towards specialized differentiated cells. The process is driven forward via the sequential activation and repression of a relatively small number of genes, frequently those encoding transcription factor (TF) proteins (*1*, *2*). By controlling the transcriptional activity of target genes, TFs establish cellular identity by determining which genes cells express. Moreover, through the modulation of various epigenetic mechanisms (e.g. posttranslational modifications of histone, DNA methylation, three-dimensional arrangement of the genome), TFs also influence the developmental potential of cells, making them more or less capable of generating individual specialized differentiated cells during subsequent steps in the process.

Current approaches for genetically programming cells attempt to override the genomically encoded stepwise progression through intermediate states by forcibly expressing TFs important for a desired final cell state. Seminal discoveries demonstrated effectiveness of this approach for reprogramming somatic cells to induced pluripotent stem cells (iPSC) (*3*), differentiating fibroblasts to myoblasts (*4*), and reprogramming non-neural cells into neurons (*5*). More recently, large-scale searches revealed TF combinations capable of programming iPSC into a variety of other cell types, including vascular endothelial and oligodendrocyte like cells (*6*, *7*). However, by skipping intermediate stages, such approaches may limit cellular function due to persistence of epigenetic features of the starting cell type and incomplete acquisition of target cell type features (*8–10*). In addition, many of the most therapeutically impactful cell types (e.g. pancreatic beta cells, hematopoietic stem and progenitor cells) appear to require a stepwise process for their differentiation and have been unattainable with an all-at-once gene programming approach (*8*, *11*, *12*).

Synthetic biology solutions intending to overcome such limitations typically employ molecular gates that can activate TFs and other genes in multiple, sequentially-delimited steps. By combining efficient and functional individual molecular gates, synthetic gene circuits could be constructed in accordance with progression through intermediate states observed in normal cell lineages (*12*, *13*). To be effective, the core molecular gates and the synthetic circuits must be competent for several essential parameters: gates must be specific by remaining inactive until triggered, gates must efficaciously activate genes once triggered, and circuits must be efficient in terms of the percentage of cells that execute a sequence of steps after being triggered.

RNA-guided nucleases and CRISPR components are attractive tools for engineering mammalian synthetic biology systems because of their inherent biochemical simplicity, specificity of DNA binding, and programmability for addressing nearly any genomic locus (*14*). Previous studies have developed methods enabling spatiotemporal regulation of spCas9 (*15–17*). Although regulation of the nuclease provides control over all Cas9-activities, it does not enable modular control at a per-gene level, which is mediated by the guide RNA molecule bound to Cas9 (*18*). Multiple studies have generated activatable guide RNA by manipulating the conformation of RNA with addition of toehold base-pairing sequences or aptamers controlled by small molecules or light (*19–22*). Unfortunately, none of the existing systems have achieved requisite levels of specificity, efficacy, efficiency, or complexity to execute more than a two-step program in cells.

In contrast to systems using conformational changes to modify guide RNA molecules, the previously described proGuide system utilized Cas9-mediated editing of a DNA encoding an inactivated guide RNA as the principal molecular gate (*23*). Conceptually, separation of the triggering mechanism (DNA editing) from the activity of the molecular gate (guide RNA) provided advantages for achieving competence in the essential parameters described above. In practice, several characteristics of the proGuide system precluded it from being readily used for building synthetic circuits for programming cell lineage progression. To meet the demands for cell programming, we systematically identified causes underlying inefficiencies in proGuides and adapted the system to result in transcriptional regulation of endogenous genes. The process resulted in a highly efficient molecular gate unit, which formed the basis of synthetic gene circuits that activated multiple genes in preprogrammed sequential orders events in individual cells. The new proGuide-based system satisfied essential parameters described above while requiring no genetic engineering of mutation of the genome either in the starting cells or throughout their use. We suggest that the proGuide system provides a new tool for incorporating rich understanding of sequential gene regulation to engineer synthetic biology solutions to cell differentiation.

## Results

### RNA Pol III terminator signals inactivate proGuides without the need for cis-acting ribozymes

The proGuide was considered for its ability to function as the molecular gate for construction of genetic circuits to program multiple, stepwise activation of endogenous genes in mammalian cells. Cas9-mediated removal of the inactivation moiety from the proGuide plasmid DNA converts the guide RNA template to encode a functional guide RNA called a matureGuide (*23*). By matching the target sequence encoded by the 20 nt spacer region from one guide RNA (**Fig. 1A**, black circles) to the 23 bp Cas9 target sequences (CTS) embedded within another proGuide’s DNA (**Fig. 1A**; white circles), a cascade of several proGuides can be activated in a sequential manner. As a cascading system, inefficiencies at each upstream step affect all downstream steps such that a relatively low level of non-specific activity (i.e. leak) in the absence of a trigger is compounded at each step disrupting the syntax of the system. Ideally, there would be no leak from each individual proGuide in the absence of a trigger. In terms of level of activity when a trigger is present, we suggest that an optimal proGuide should display the same activity in cells as an sgRNA does. Using these performance targets, previously published proGuides (v1.0)(*23*) with insertions in the tetraloop or hairpin 1 location were significantly less effective than an sgRNA for disruption of EGFP, and the tetraloop variant displayed significant leak in the absence of a trigger guide RNA (**Fig. 1B**).

**Fig. 1.**
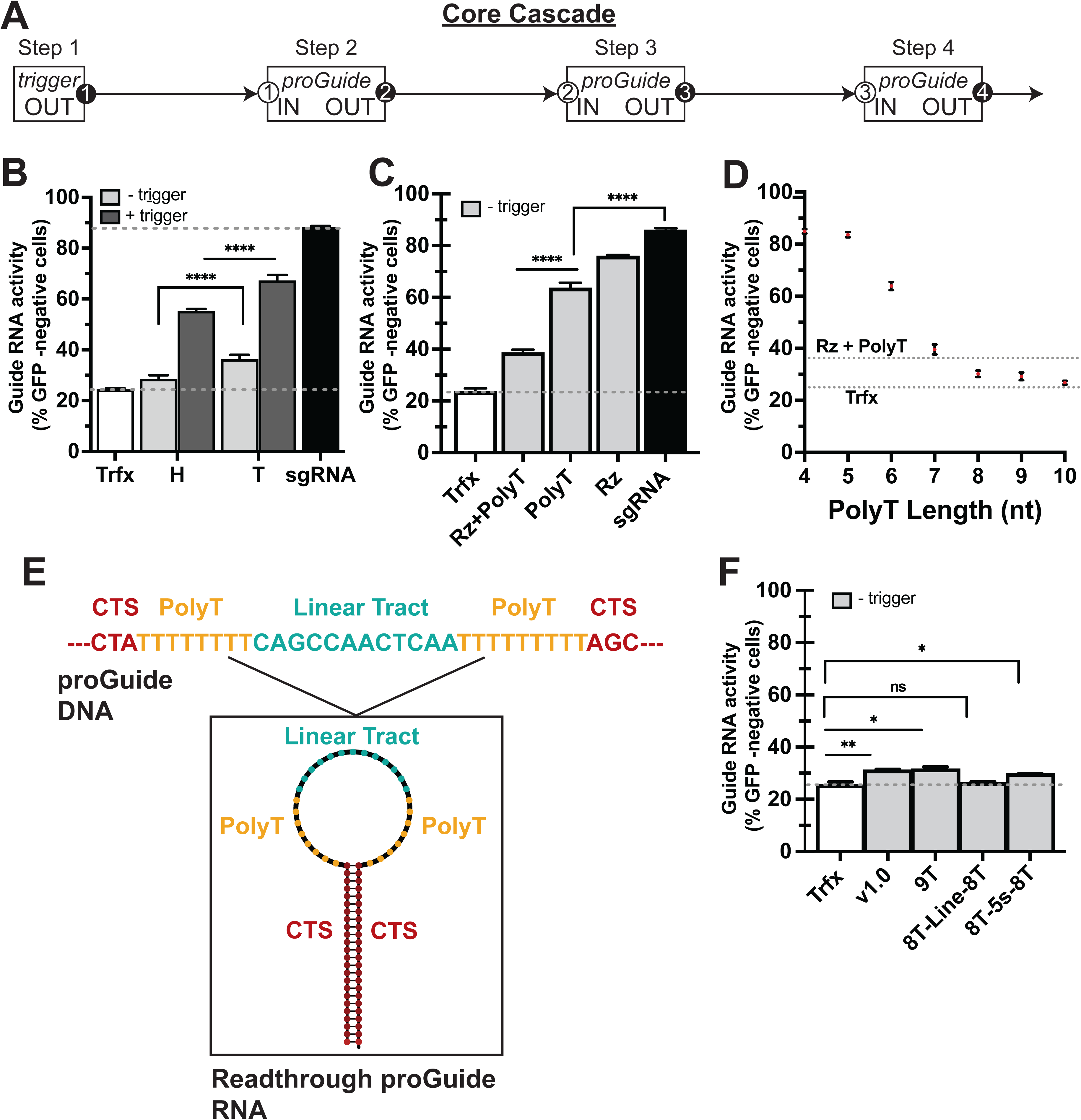
Inactivation proGuides by RNA Pol III transcriptional termination sequences. **A**) Schematic of a cascade of proGuide plasmids wherein the spacer sequence functions as an output activity (black oval) of one guide and is matched to the Cas9 target sequences (CTS; white oval) as the sensing input present in a downstream proGuide DNA. **B-D**) Reduction of GFP-positive HEK293T cells harboring a genomic EGFP expression cassette was determined by flow cytometry 72 h after transient transfection of guide RNA plasmids and a Cas9 expression plasmid. Control transfections either lacked an EGFP targeting guide RNA (Trfx; white bars, dotted line) or included an sgRNA targeting EGFP (sgRNA; black bars). **B**) First generation proGuides displayed activity in the absence of a trigger RNA (light gray bars) when the inactivation sequences were located in the tetraloop (T) region and incomplete conversion to an active state when located in the hairpin (H) region. **C**) Inactivation with only the ribozyme (Rz), only the 6nt polyT tract (PolyT) or both (Rz+PolyT) all displayed leakiness. **D**) Increasing the length of the polyT tract reduced leakiness of EGFP proGuides below that of the first generation inactivation sequences (Rz+PolyT dotted line). **E**) Schematic of the DNA sequence (top) encoding a dual polyT tract and predicted RNA secondary structure (bottom). **F**) Absence of detectable leak from proGuides harboring the dual polyT tract depicted in (E). All data represent mean of biological triplicates +/- standard deviation. * indicated p<0.001; ns indicates p>0.05.

To elucidate the causes of deficiencies and to improve system performance, proGuide leak and activity were tested independently. Initial proGuide designs had incorporated redundant mechanisms in the DNA sequence designed to disrupt the activity of the encoded guide RNA (*23*). A cis-cleaving hammerhead ribozyme sequence was included to cleave the RNA prior to binding to Cas9, and a polyT tract of six consecutive thymidine residues was designed to terminate RNA transcription preventing synthesis of a full-length functional guide RNA (**Fig. S1**). The impact of each inactivation moiety was tested by constructing proGuide variants possessing either a hammerhead ribozyme or the polyT tract alone within the loop structures of the tetraloop or hairpin 1. The polyT tract alone was more effective than the ribozyme alone in mitigating unwanted guide RNA activity when placed in either the tetraloop or the hairpin locations (**Fig. 1C, Fig. S2**). Conversely, using a full-length hammerhead ribozyme with purported enhanced RNA nucleolytic activities as the inactivation moiety unexpectedly exacerbated proGuide leak (**Fig. S3**). Given that the enhanced activities of the full-length ribozyme are dependent on RNA folding (*24*) and placement of the ribozyme within the context of a proGuide may compromise this folding, it could be challenging to design a system of leak-free proGuides using ribozyme-based inactivation strategies.

Several previous observations supported focusing on the polyT tract for development of more effective inactivation moieties. Inactivation by the polyT tract relies on its ability to terminate elongation of RNA polymerase III (RNA Pol III), precluding transcription of the downstream gRNA sequence, which is necessary for Cas9 activities (*25*). Several known parameters, including the number of consecutive thymidines, the promoter initiating transcription, and the nucleotide sequence upstream of the polyT tract have been shown to control the extent of termination (*26*, *27*). When incorporated as the sole inactivation moiety in proGuides, increasing the number of consecutive thymidines in a polyT tract progressively reduced leak, with the most substantial mitigation in leak observed when lengthening the polyT tract from four to eight thymidines (**Fig. 1D**). Addition of a 9^th^ or 10^th^ consecutive thymidine provided minimal additional reduction in activity (**Fig. 1D, Fig S4**). DNA encoding the RNA fragment expected from termination by the polyT tract and lacking guide sequences downstream of the polyT tract failed to disrupt EGFP in cells (**Fig S5**), indicating that the residual leak resulted from transcriptional readthrough rather than a low-level activity of the RNA fragment terminated at the polyT tract.

To eliminate the residual leak, a dual polyT tract strategy similar to those observed in some tRNA genes was adopted (*26*). Given that the RNA secondary structure preceding the terminator has been suggested to enhance termination (*27*), two different DNA sequences were evaluated for the RNA structures that they encoded. One sequence encoded a 5S rRNA sequence that was previously shown to stimulate RNA Pol III termination in mammalian cells, whereas the other encoded an RNA motif predicted to confer a linear conformation (*27*). Both structures were flanked between the two terminator tracts (**Fig. 1E**). Surprisingly, configuration of two 8T tracts separated by the linear RNA conformation resulted in the lowest amount of leak, which was comparable to the control condition without any guide RNA (**Fig. 1F**). Given that these results were from tetraloop-modified proGuides, which had previously displayed greater leak and higher conversion activity (*23*), this inactivation sequence was used as the foundation for further development of the proGuides.

### Orientation of the Cas9 target sequences in the proGuide DNA template determines DNA repair and efficacy of the matureGuide

The efficacy of a proGuide as a molecular gate in mammalian cells depends on both the frequency of its conversion to a functional matureGuide and the activity of the matureGuide RNA encoded by the edited DNA template. Ideally, a matureGuide would exhibit the same level of activity as an sgRNA; however, previously published matureGuides frequently displayed activity significantly lower than that of an sgRNA (**Fig. 1B, S2**) (*23*). Although the Cas9 sgRNA has been amenable to additions of sequence in the loops of the tetraloop and hairpin 1, even minor modifications to the sgRNA structure can affect guide RNA activity (*25*, *28*), suggesting an area to evaluate for improving proGuides.

To determine if the proGuide architecture could affect the activity of matureGuides, different proGuide configurations were made and assessed for activity in response to a trigger. The CTS sequences were identical for all proGuide variants to eliminate potential confounding effects of variations in EGFP disruption caused by the activity of the trigger sgRNA on the CTS. Similarly, the inactivating moiety was also kept constant in the five proGuide configurations, which differed by orientation of the CTS as a direct repeat (DR) versus inverted repeat (IR), by the presence of additional DNA repeats nested between the two CTS, and by the presence of additional DNA repeats flanking the two CTS (**Fig 2A**). Secondary structures of the RNA sequence at the end of the tetraloop were predicted assuming perfect non-homologous end joining (NHEJ) repair of the DNA (**Fig 2A**), and they illustrated the base-pairing of nucleotides encoded by inverted repeats forming additions to the stem structure in the tetraloop region (**Fig. 2A**).

**Fig. 2.**
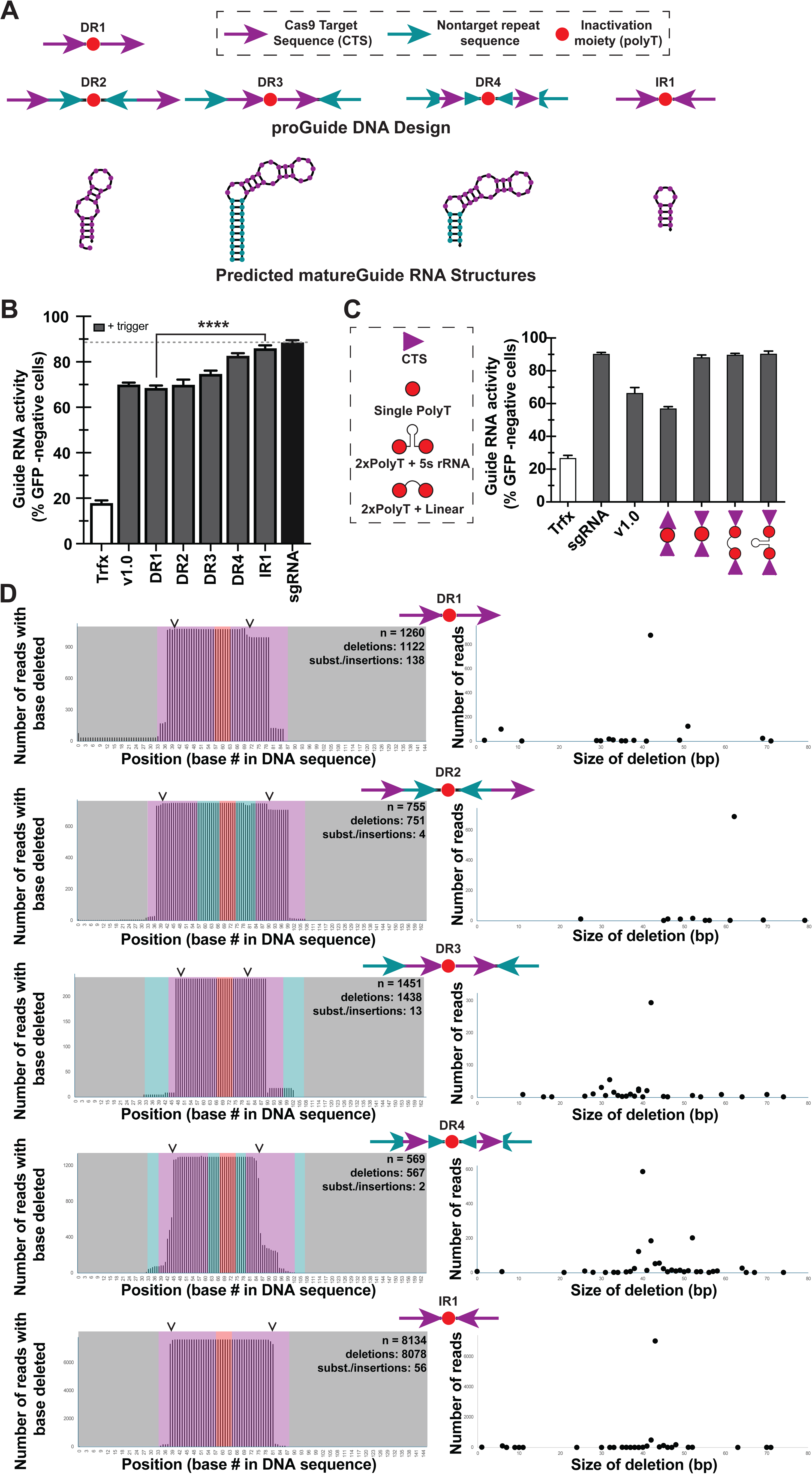
Effects of the orientation of CTS repeats on conversion of proGuides to active guide RNA. **A**) Schematic of the arrangement of different proGuide components in a DNA sequence (top) and predicted RNA sequence structures (bottom) following a perfect NHEJ repair-mediated nested deletion between the two Cas9 cut sites. **B**) Transient transfections (as described in Fig. 1) with proGuide expression plasmids depicted in (A). **C**) Conversion of IR1 configuration to an active guide RNA state was not significantly changed by the termination sequences (schematic on left). **D**) DNA sequencing results of reverse transcribed guide RNA from transiently transfected cells were analyzed based on the frequency of different sizes of RNA mapped to proGuide sequences. The frequency of deletions at each base in the proGuide (left) and the distribution of sizes of deletions (right) for each proGuide with the configurations shown in (A). Data represent all sequences with at least two reads in a sample.

In cells, these new configurations displayed a range of matureGuide activities (**Fig 2B, Fig S6**). Although DR1 and DR2 variants displayed better inactivation of the proGuide (**Fig. 1C**) and had a simpler design relative to previously published proGuides (v1.0), these new variation did not improve editing activity in response to a trigger (**Fig. 2B**). Placing DNA repeat sequences to flank the CTS in DR3 and DR4 increased activity in response to a trigger, suggesting that the stem structure resulting from the repeat sequence supported functionality of the matureGuide RNA (**Fig. 2A,B**). Interestingly, the shorter stem structure of DR4 resulted in higher matureGuide RNA activity than the longer stem from DR3. Consistent with a role of the stem structure for activity of matureGuides, the IR1 configuration generated the highest matureGuide RNA activity. Note that perfect NHEJ repair between two CTS configured in a so-called PAM-out orientation of IR1 results in 12 bp inverted repeat consisting of the 3 bp PAM sequence and PAM-proximal 3 bp from each CTS. The predicted post-NHEJ structure for the IR1 configuration had one of the most stable hairpin structures, which would support the tetraloop structure of the guide RNA. Replacing the inactivation moiety in these variants with more effective ones (**Fig. 1D, E**) indicated that effectiveness of the inverted repeat orientation was not dependent upon a specific sequence between repeats (**Fig 2C**). The superiority of inverted repeat orientation was also consistent in proGuides with other CTS sequences (**Fig S7**), suggesting the effect was not caused by altering the nuclease activity of Cas9 at individual CTS sequences. Since previous research also demonstrated increased guide RNA activity from stabilization of the tetraloop (*28*), we suggest that the six bp supporting stem structure from IR1 configurations likely stabilized the hairpin structure of the matureGuide tetraloop, improving efficacy of matureGuides.

To determine the effect of DNA repair on the efficiency of proGuide conversion to matureGuides, we began with an RNA sequencing approach. Briefly, cells were transfected with proGuide, Cas9, and trigger sgRNA plasmids, and RNA was isolated and amplified for sequencing of RNA molecules possessing the 3’ guide RNA sequence, i.e. downstream of the polyT tract. Cas9-mediated changes to proGuide plasmids were characterized by mapping sequencing reads onto the proGuide DNA reference sequence. This RNA sequencing approach showed that the previously published proGuide architecture generated a diverse set of RNA sizes in cells (**Fig. S8**), including 35% of sequences aligning to the 254 bp proGuide sequencing caused by readthrough transcription and failure of ribozyme cleavage of the proGuide RNA transcript. Surprisingly, 166 bp RNA, corresponding to the size expected from perfect NHEJ repair, made up only 1.5% of transcripts (**Fig. S8**). In contrast, conversion of proGuides harboring CTS in an inverted repeat orientation produced RNA sequences corresponding to the perfect NHEJ repair between the two CTS (**Fig. 2E, Fig. S8**). Comparison of RNA produced from conversion of DR1-4 and IR1 proGuides (**Fig. 2A,B**) revealed none of the DR configurations generated RNA corresponding to perfect repair between CTS (**Fig. 2D, Table S2-S7**). Instead, most frequent outcomes had additional deletion of CTS sequences, both upstream and downstream of the Cas9 cut site (arrowheads in **Fig. 2D**). The DR4 configuration produced the most variable outcomes with the predominant 40 bp loss occurring in only 40% of deletions shown in Fig. 2D. Outcomes from conversion of DR1 were notable, because it was the only configurations to produce a substantial frequency of non-deletion changes, which included 5% of mapped reads corresponding to a perfect inversion of the sequence between the CTS (**Table S2**). By contrast to the DR configurations, the majority (63%) of outcomes from IR1 corresponded to a perfect deletion between the two CTS (**Fig. 2D, Table S2**). Regardless of CTS configuration, less than 1% of RNA sequences corresponded to readthrough guide RNA, which was consistent with very low levels of leak activity from this proGuide in cells (**Fig. 1E**). The frequency of RNA corresponding to either a perfect repair or insertion/deletion of 1 bp between the two CTS from IR1 was consistent with structurally permissive and thus highly effective form of matureGuide RNA displaying high activity in cells. The dramatically different frequencies of RNA sequences produced by proGuides with the same CTS sequences, but in different orientations, suggests that the DNA repair process is affected by CTS orientation. One possible explanation for failure of DR1-4 is that perfect NHEJ repair of direct repeat CTS would form an intact single CTS in the plasmid, which would provide a substrate for additional Cas9 cutting.

### High efficiency CTS sequences are required for multi-step cascades of proGuides

Whereas experiments evaluating the inactivation moiety (**Fig. 1**) and the proGuide architecture (**Fig. 2**) kept the CTS sequence constant to control for effects of Cas9 nuclease activity, we anticipated that CTS sequences would require optimization to build efficient cascades of multiple proGuides. We considered that one potential limitation could come from competition for common resources (e.g. Cas9-VPR protein, number of transfected plasmids, RNA Pol III, etc), and maintaining efficiency of each individual proGuide may be challenged in cells loaded with multiple proGuide plasmids and matureGuide RNA. To determine effects of the CTS sequences when challenged with competition, EGFP disruption was measured in cells transfected with different proGuide:trigger sgRNA ratios, ranging from 20:1 to 0.05:1 (**Fig. 3A**). Regardless of the relative amount of the CTS101-containing proGuide, it disrupted the genomic EGFP gene similarly to the sgRNA control, and the proGuide was notably effective even at lowest concentrations representing 2% of total plasmid DNA delivered to cells. The CTS102-containing proGuide generated the lowest levels of EGFP disruption at all concentrations (**Fig 3A**), suggesting a deficiency in the matureGuide RNA activity independent of activity of the CTS102-specific trigger sgRNA. The CTS103-containing proGuide displayed sgRNA-like activity at low proGuide concentrations and reduced activity when the relative amount of trigger RNA was decreased, suggesting a deficiency in conversion by the CTS103-specific trigger sgRNA (**Fig. 3A**).

**Fig. 3.**
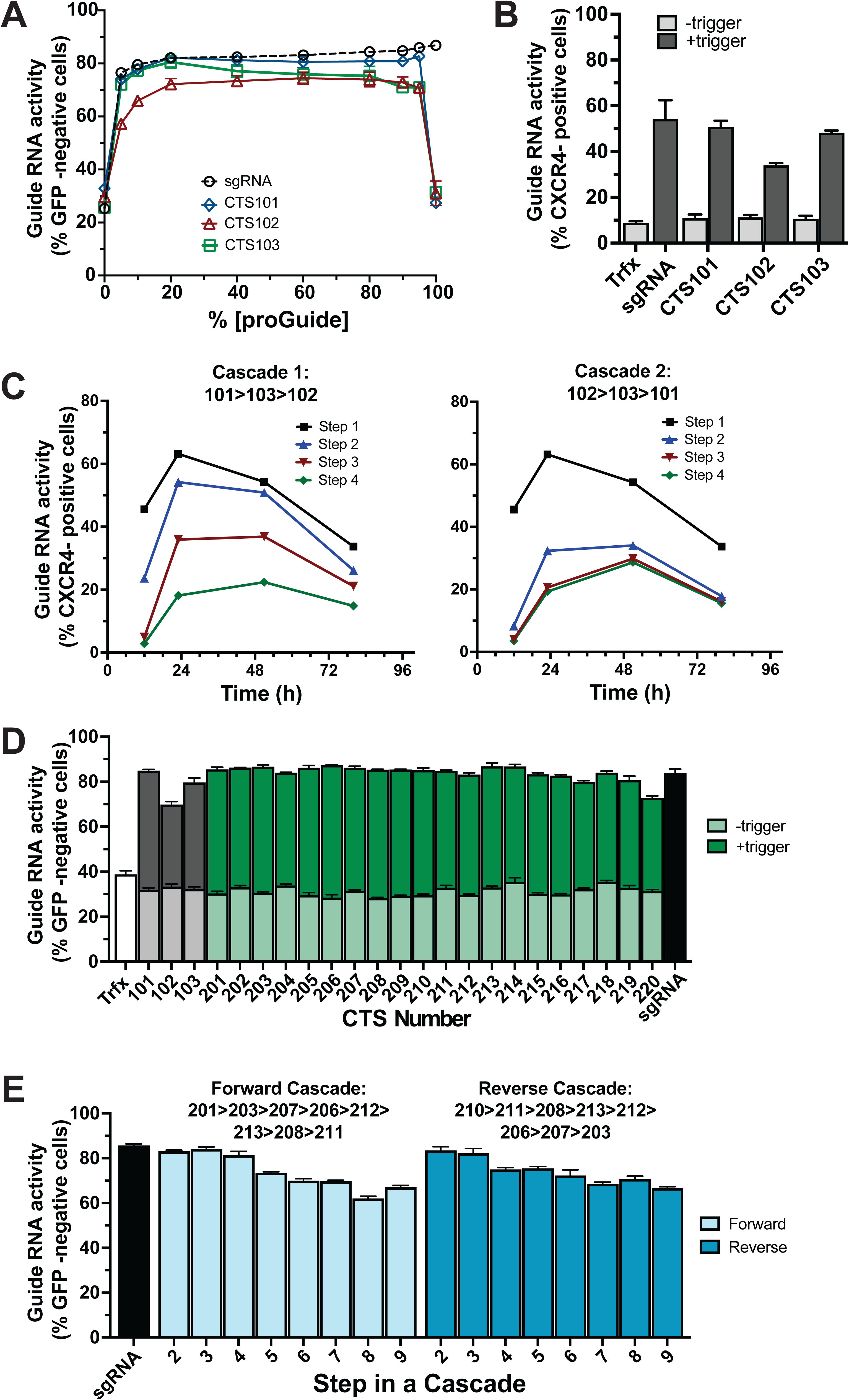
Sequence composition of CTS affects efficiency of proGuide conversion. **A**) The EGFP disruption activity of different ratios of proGuide : trigger plasmid DNA was determined by flow cytometry 72h after transient transfection (similar to Fig. 1). Three proGuides and their matched trigger plasmids used different CTS (CTS101, CTS102, CTS103). **B**) Activation of the endogenous CXCR4 gene determined by cell surface protein expression was determined by flow cytometry 48 h after transient transfection of a proGuide (CTS101, CTS102, CTS103) with a 14nt spacer targeting the CXCR4 promoter, a trigger plasmid, and a Cas9-VPR expression plasmid. Transfections lacking a guide RNA and with an sgRNA with the 14nt CXCR4 spacer were used as negative/positive controls. **C**) Two cascades of proGuides constructed using spacers and CTS sequences corresponding to CTS101, CTS102, and CTS103 (Fig. S10), were transiently transfected into HEK293T cells with a Cas9-VPR expression plasmid, and assessed by flow cytometry for surface protein expression of CXCR4. **D**) Activity of proGuides (+/- trigger) harboring 20 new CTS sequences determined by EGFP disruption (similar to as described in Fig. 1). **E**) Two cascades constructed of proGuides using new CTS sequences (D) in the sequential order indicated above the graph.

To evaluate effects of CTS sequences on proGuides for transcriptional activation of endogenous genes, a nuclease active Cas9-VPR fusion protein was used for both conversion of proGuides to matureGuides and for CRISPRa activation of genes. The two activities can be separately targeted by using 20nt spacers for Cas9 nuclease activity and 14nt spacers for DNA binding to promoters of target genes (*29*). The CXCR4 cell surface protein was used as an endogenous reporter gene due to the simplicity of detection by flow cytometry and previous reports showing its activation by Cas9-VPR (*29*). Several 14 nt spacer sequences were evaluated for activation of CXCR4 expression in HEK293T cells (**Fig. S9**), and the most active spacer was cloned into proGuide plasmids. Plasmids encoding a proGuide with a 14 nt spacer targeted to CXCR4, a trigger sgRNA, and Cas9-VPR expression were co-transfected into HEK293T cells and the frequency of cells displaying CXCR4 expression was measured after 48 hours by flow cytometry (**Fig. 3B**). Given that CXCR4 expression was dependent upon the inclusion of a trigger sgRNA plasmid, these results indicate that Cas9-VPR was able to catalyze both the conversion of a proGuide to a matureGuide and the activation of CXCR4 transcription from its endogenous gene. Thus, a proGuide-based system can be used to activate endougenous genes via the expression of a single Cas9 fusion protein. The different frequencies of CXCR4 activation from proGuides with different CTSs suggests that the activation of an endogenous gene may be more sensitive than the disruption of EGFP to conversion efficiency and efficacy of matureGuides.

Effects of CTS sequences with variable activities were tested by assembling multistep proGuide cascades, which required Cas9 nuclease for stepwise progression and required VPR-stimulated transcriptional activation for endogenous gene expression (**Fig. S10**). Two cascades were generated from proGuide plasmids with spacers from one step designed to target the CTS in the next step’s proGuide (**Fig. 3C, S10**). Changes to the cell surface expression of CXCR4 over time indicated that proGuides containing CTS102 displayed reduced activation substantially at the step requiring CTS102 and also at all downstream steps (**Fig. 3C**). CTS103 also caused reduced activation in both cascades, and due to the successive nature of cascades, this resulted in a low-level activation of steps 2, 3 and 4 in cascade 2. By contrast, steps requiring proGuides with CTS101 displayed the highest activity compared to previous steps. Interestingly, the inefficiency of CTS102 combined with high efficiency of CTS101 resulted in a timing defect in cascade #2, whereby step 3 (CTS102 proGuide) and step 4 (CTS101 proGuide) failed to display the syntactic separation expected from the design of the cascade. Together, these results identified possible modes of failure for cascades of proGuides and suggested a need for new CTS sequences to generate efficient and effective proGuides.

To generate new sequences to function as CTS in proGuides, a virtual library of two million random 23 bp CTS sequences were generated and filtered to remove sequences with features (e.g. GC content, PAM-distal guanines, PAM-proximal thymidines), previously shown to impair Cas9 activity (**Fig. S11**)(*30–33*). To minimize potential Cas9 nuclease activity on the human genome and other commonly used animal genomes, CTS sequences were filtered to ensure at least seven mismatches with any sequence in genome assemblies from human, mouse, cow, pig, tuna, and chicken. Restriction through filters resulted in over 50,000 possible CTS sequences, all predicted to lack on-target and off-target Cas9 activity for the genomes of the selected organisms (**Fig S11**).

A set of 96 of these CTS were randomly selected for empirical testing in human cells. The selected CTS and the matching spacer sequence were cloned into a plasmid enabling the expression of an EGFP-targeting proGuide and its trigger sgRNA from a single plasmid. Loss of GFP fluorescence 48 hours post-transfection showed a range of CTS functionality, including several as active as CTS101 (**Fig 2E, Fig S12**). The top twenty CTS sequences were cloned into separate proGuide plasmids, their matching spacers were cloned into trigger sgRNA expression plasmids, and each combination’s ability to disrupt EGFP in cells was compared to CTS101-103 proGuides (**Fig. 3D**). As expected, twenty proGuides containing CTS201-220 sequences exhibited negligible leak and high matureGuide activity when delivered with its trigger sgRNA on a separate plasmid (**Fig. 3D**). Next, new proGuide plasmids containing CTS201-220 were constructed with the 14 nt spacer for transcriptional activation of CXCR4 and evaluated for activity with Cas9-VPR. In this CRISPRa activity assay, all proGuides displayed negligible leak and guide to guide variability for matureGuide activity (**Fig. S13**). Several CTS (CTS201, 203, 207, 206, 212, 213, 208, 211) that performed well in both the EGFP disruption and CXCR4 activation assays were chosen for evaluation in new proGuide cascades using only these top performing CTS2xx generation of sequences.

Initial tests of new CTS in proGuide cascades measured the ability of an EGFP-targeting proGuide placed at each individual step in 9-step cascades. Each 9-step core cascade was generated by inserting 20 nt spacer sequences into proGuide base plasmids such that each proGuide would target the CTS of the next proGuide in either forward or reverse cascades depicted atop **Fig. 3E**. Each transfection mix contained proGuide cascade plasmids, a Cas9-VPR expression plasmid, and one EGFP-targeting proGuide that possessed the CTS programming activation at only one step from 2 to 9. In cells, the overall frequency of EGFP disruption after 72 h decreased slightly with the progression of each step in both the forward and the reverse cascade (**Fig. 3E**). Notably, the stepwise decrease in activity did not correspond to specific CTS, suggesting a high degree of modularity with the chosen proGuide CTS and spacer components.

### proGuide cascades enable sequential and multiplexed transcriptional activation of endogenous genes

To determine whether a cascade of proGuides could promote stepwise activation of endogenous genes, a core cascade of five proGuides was combined with a proGuide targeting CXCR4 for transcriptional activation at one step based on the identity of its CTS. Similar to proGuide cascades used for disruption of EGFP (**Fig. 3E**), the CXCR4-activation cascades were efficient in terms of the percent of cells that completed a 6-step cascade and activated the last proGuide step (**Fig. 4A**). Comparison of cascades at different time points showed that placement of the CXCR4 proGuide at different steps resulted in the expected sequential activation of CXCR4 expression, with a lag period of about 6 hours between steps (**Fig. 4B**). Experiments examining all combinations of cascades with proGuides containing CTS201, 203, and 207 for steps 2, 3, and 4 indicated that these different proGuides functioned similarly within cascades (**Fig. 4C**), supporting the modularity of proGuides as a molecular gate for the activation of endogenous genes by CRISPRa. Additionally, 10-step cascades were evaluated to test the limit to the number of steps that could be programmed with the CTS2xx-containing proGuide plasmid DNAs (**Fig. S14**). The kinetics of CXCR4 activation in these 10-step cascades displayed sequentially separated activation up until step 8, at which point the syntactic separation distinguishing activation of individual steps was lost (**Fig. S14**).

**Fig. 4.**
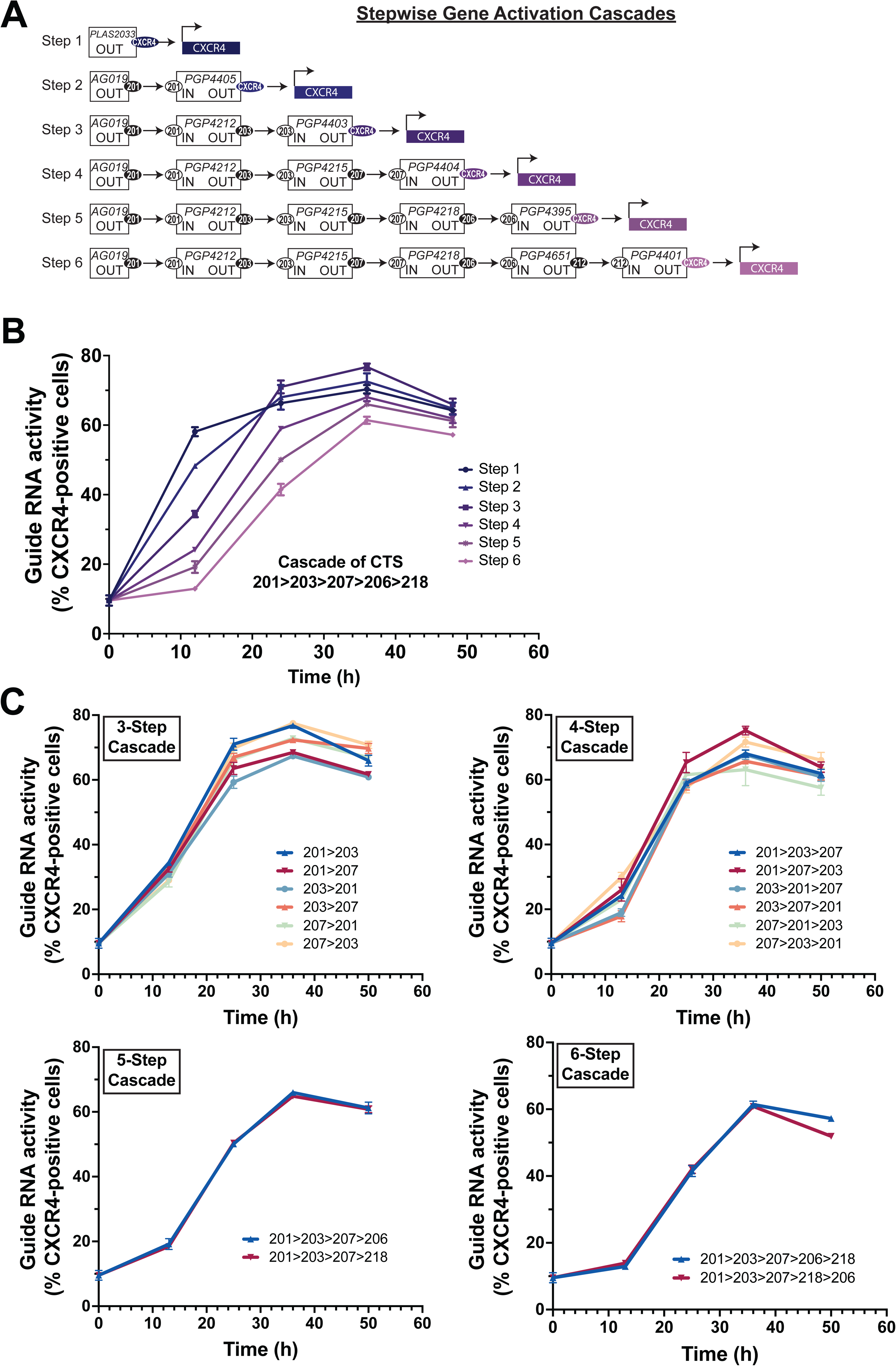
Control over kinetics of endogenous gene activation by transient transfection of proGuide plasmids. **A**) Schematic depicting plasmid DNA composition of proGuide cascades used in (B), as described in (Fig. S10). **B**) CXCR4 surface protein expression was measured by flow cytometry at the indicated times after transient transfection of HEK293T cells with Cas9-VPR and plasmid mixes depicted in (A). **C**) The order of the CTS used in cascades of proGuides was iterated on by generating new proGuide plasmids for step wise progression to activation of CXCR4 transcriptional activation by Cas9-VPR. Each graph shows CXCR4 surface protein expression for cascades designed to activate CXCR4 at the indicated step number.

To direct cellular differentiation through steps in a lineage, proGuide cascades need to be able to activate multiple endogenous genes in a preprogrammed syntactic manner. We began to evaluate such capabilities in HEK293T cells by using proGuides to program the activation of two distinct genes (CXCR4 and CD105) at different steps in cascades of proGuides (**Fig. 5A**). Briefly, a 7-step core cascade was used for all conditions, and proGuides activating the endogenous genes were included for activation in steps 1, 4 and 7. Kinetics of activation were examined by flow cytometry every 12 hours after DNA delivery. In transfections where both genes were programmed for activation at step 1 via sgRNA, cell surface expression of CD105 occurred approximately 12hrs after CXCR4, suggesting a delay in CD105 independent of any control programmed by proGuides cascades (**Fig. 5B**). Switching the order of CXCR4 and CD105 in step 1 vs 4 or step 1 vs 7 changed the order in which the surface expression of the two genes products were detected on individual cells (**Fig. 5B**). Consistent with cascades of proGuides activating expression of only one gene, placing activation proGuides at a later step in a cascade delayed gene product expression relative to earlier step activation.

**Fig. 5.**
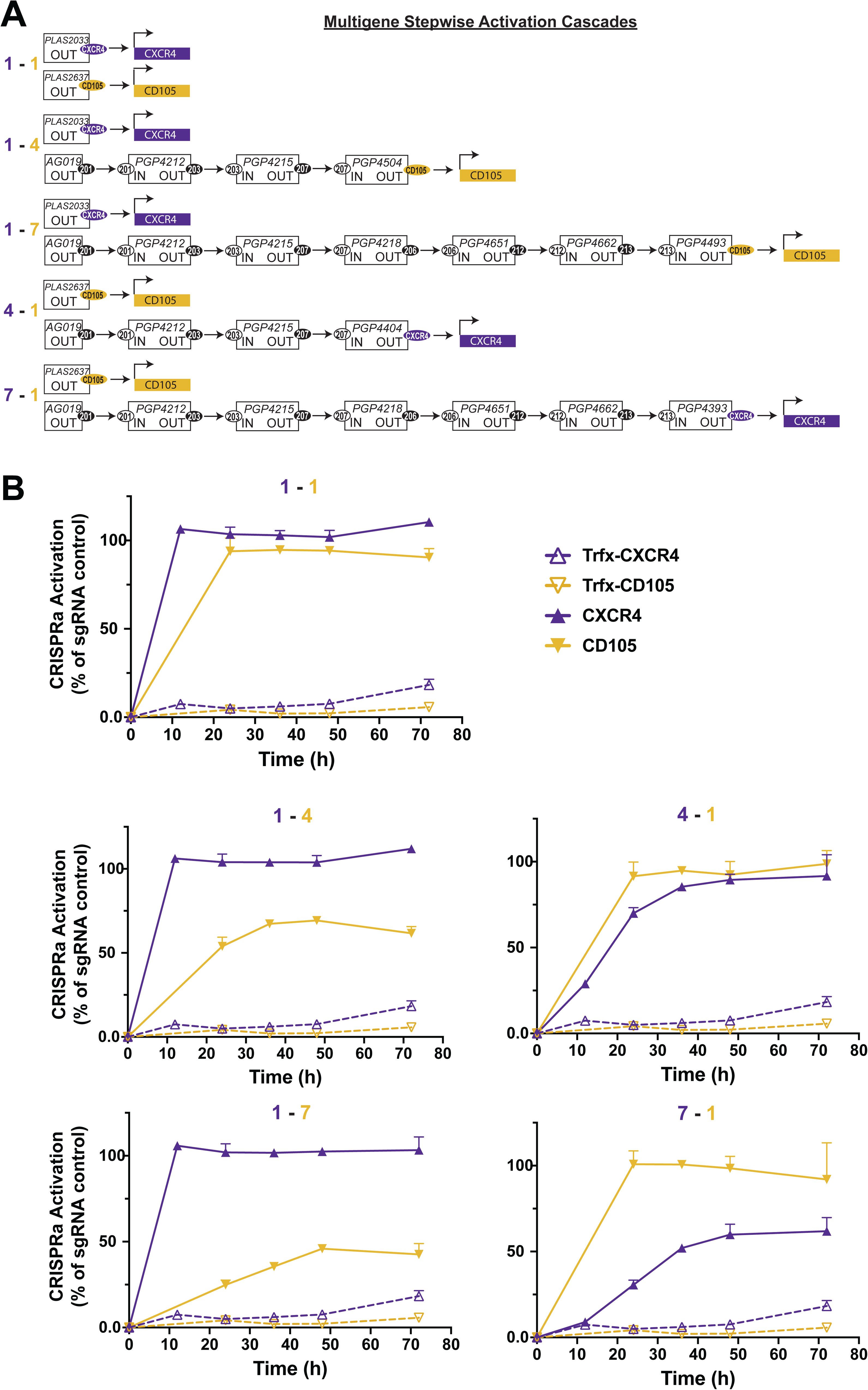
Syntactic activation of CD105 and CXCR4 genes using cascades of proGuides. **A**) Schematic depicting plasmid DNA composition of proGuide cascades used in (B). **B**) CXCR4 and CD105 surface protein expression was measured by flow cytometry at 12 h intervals after transient transfection of HEK293T cells with Cas9-VPR and plasmid mixes depicted in (A). Graphs show the percentage of cells positive for CXCR4/CD105 relative to the number of positive cells observed in sgRNA-positive control transfections at each time point. Note, that since the sgRNA positive control did not display CD105 positivity at 12 h, normalized data are not shown for CD105 at that time point. Non-normalized data are presented in Fig S13?.

Finally, to test the preprogrammed syntactic activation of several factors in individual cells, cascades of proGuides were engineered to activate endogenous genes encoding CD105, CD4, and DLL4 (**Fig. 6A**). Pools of four guide RNAs were generated for each gene based on previously reported guidelines (*34*) and were targeted to be triggered at the same step by using the same CTS in each of the four proGuides for a single CRISPRa target (**Fig. 6A**). The use of four guides for each target provided more robust gene activation (**Fig. S15**) and also provided insight into the level of complexity available designing plasmid-based cascades, as these required up to 19 different plasmids to be introduced into individual cells for a complete instruction set. The cascades were tested in human iPSC to better evaluate potential for programming cell differentiation in a stem cell system capable of making any somatic human cell type. After nucleofection of plasmid DNA, the cell surface expression of target gene products was consistent with the stepwise activation of gene products corresponding to their positions in the proGuide cascades (**Fig. 6B**). Interestingly, the different cascades revealed gene-specific effects that were not apparent when a single gene was activated at multiple steps in cascades (**Fig. 4**). Within cascades, activation of DLL4 surface expression was rapid relative to slower activation of CD4 and CD105 surface expression. Comparison of the rates of activation indicated that earlier steps (step 1, step 2) produced a more rapid increase among the population of cells than the later steps (step 5, step 7) (**Fig. 6C**). Notably, when all three targets were programmed to activate in the same cells at step 1 (**Fig. 6D**), the surface expression of DLL4 and CD4 was slower than when programmed by itself at step 1 in a cascade (**Fig. 6B,E**). Taken together, these data demonstrate capabilities of cascades of proGuides for installing diverse arrangements of sequential gene activation programs in human stem cells. They idicate an ability to turn on several genes at different times relative to one another in individual cells. Given the flexibility of Cas9-VPR for activation of virtually any gene in the genome, complex programs of sequential gene activation processes are possible.

**Fig. 6.**
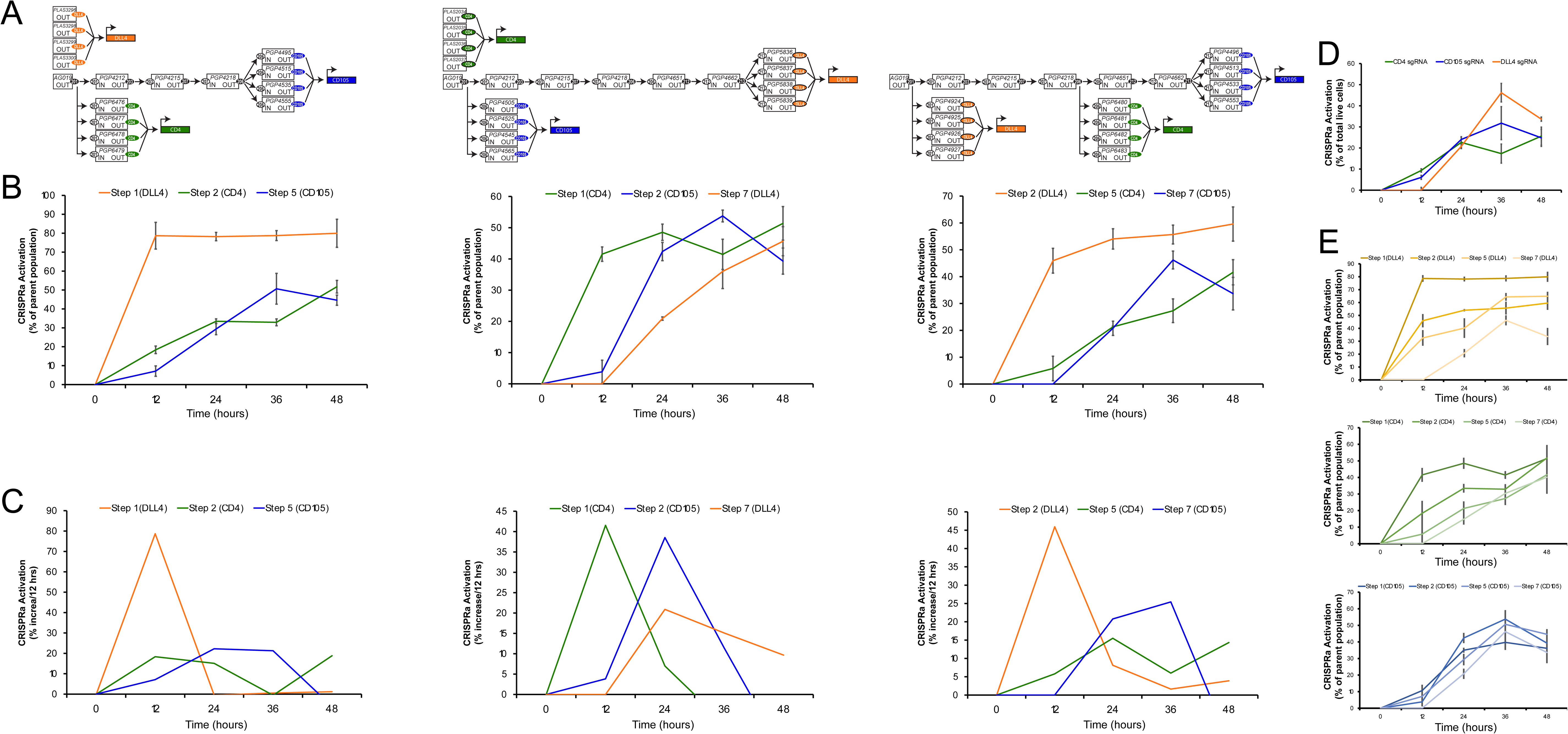
Syntactic activation of several genes using cascades of proGuides. **A**) Schematics depicting plasmid DNA composition and arrangements of proGuides for the three cascades used in (B,C). **B**) The expression of the target surface marker proteins in human iPSC after nucleofection was measured every 12 hours by flow cytometry. Values are mean +/- standard deviation of three replicates and correspond to the percentage of cells expressing the indicated marker from the pool of cells that expressed the marker(s) upstream in the cascades. **C)** The rate of activation of each gene was determined from the frequencies in (B) and represented as the percent increase per 12 hours. D) CRISPRa activation of the three gene products (CD4, CD105, DLL4) in iPSC after nucleofection of pools of sgRNA for all three genes. **E)** Relative activation of surface marker proteins in iPSC where it was programmed to be activated at the indicated step by cascades of proGuides.

## Discussion

Since the discovery of the ability of MyoD to program fibroblasts to differentiate into skeletal muscle cells (*4*), significant efforts identified key TF genes and cell types amenable for programming and improved exogenous, transgenic expression of TFs in cells (*2*). Subsequently, the development of genetic engineering and synthetic biology tools have allowed more sophisticated programming using molecular gates, which can elicit cellular effects depending upon the presence or absence of triggering conditions (*35*, *36*). The work presented here advances cell programming capabilities with a system that can deploy seven sequential molecular gates, each triggered by the completion of the previous step, and each capable of activating the expression of different endogenous genes. The number of steps, the capacity to activate any gene in the genome, the ability to execute a complete gene circuit in human iPSC without genome modification, and the efficiency of stepwise progression are each significant advances in mammalian synthetic biology.

Although the focus of the current study was to transform the proGuide into an effective and modular molecular gate for programming gene activation, several findings from this endeavor could have significant and broader implications independent of proGuides. Notably, simple inactivating moieties comprised of polyT tracts decisively prevented RNA Pol III synthesis of functional guide RNA transcripts and were superior to ribozymes, likely due to the dependency upon ribozyme folding for its inactivation mechanism (*37*). The mechanism of a polyT tract- inactivation should apply to any RNA produced by RNA Pol III (*26*), assuming the polyT termination tract precedes the functional component of the RNA. As such, the polyT tract could inactivate microRNAs, small hairpin RNAs, ribozymes, and other RNA-guided enzymes, all of which typically use RNA Pol III for expression in mammalian cells. Conversion of such RNAs to an active state would depend upon the mechanism used to remove or disrupt the DNA encoding the polyT tract, as is the case for deletion via Cas9 cutting at CTS flanking the polyT tract. We suggest that the removal of polyT tracts via Cas9-deletion could provide conditionality to RNA Pol III transcripts analogous to how Lox-Stop-Lox cassettes have been used for RNA Pol II transcripts (*38*, *39*).

Analysis of guide RNA transcripts from plasmid DNA provides insights into the repair of extragenomic DNA following Cas9 cleavage in mammalian cells. In comparison to the abundance of research into the repair of genomic DNA following Cas9 cutting (*40–42*), effects on plasmid DNA are relatively unknown. Plasmid DNA harboring CTS arranged in a direct repeat configuration rarely resulted in an NHEJ repair process with a perfect nested deletion between cut sites. By comparison, repair of a genomically integrated copy of the DNA sequence resulted in nested deletion between sites in about one third of edited DNAs (*23*). Interestingly, changing the configuration of the CTS from DR to IR in plasmid DNA drastically increased the frequency of perfect nested deletions. We suggest that the perfect repair with IR configurations is consistent with frequent monomolecular (i.e. the ends of one plasmid DNA) repair process and infrequent interactions between plasmid DNA molecules. In addition, the lack of perfect repair from DR configurations can be explained by a perfect repair between the CTS resulting in a re-formation of an intact CTS, which can be cut again by Cas9. The predominant outcome from these secondary cuts appeared to result in small deletions. An exception to this possibility was the inversion of the complete DNA sequence nested between cut sites, which was observed at unexpected high frequency (∼4%) with the DR1 configuration. MatureGuides resulting from such inversion repair outcomes would feature a large bulge on the tetraloop and lack any hairpin extension that could destabilize the tetraloop, which could explain the relatively low matureGuide activity from of this configuration. The molecular mechanisms responsible for this high rate of inversion remain unclear. Together, these results suggest that repair of DSB of plasmid DNA may be affected by different parameters than genomic DNA.

The proGuide-based cascades developed here have generated new capabilities for programming cells. One distinguishing characteristic of these cascades from previously developed systems is the autonomous nature of proGuide cascades. Once they are initiated by a trigger guide RNA, the cascade progresses without the need for conditional activation from the user via an external stimulus (e.g. small molecule, light, temperature)(*15*, *20*, *43–45*) or via cellular activity (expression from conditionally active promoter, processing by a conditionally active enzyme)(*21*, *46*). Some previous systems have been developed for autonomous progression, including the initial proGuide and a system using the conversion of an inactive Cre-Lox for the inactivation moiety in guide RNA (*23*, *47*). Each of these possessed the ability to address the genome with a variety of Cas9-mediated activities similar to the system developed in the current study; however, their efficiency was extremely low, enabling only one to two steps to be completed in cells (*23*, *47*). Other systems using base editors to make changes to the sequence of guide RNA displayed capabilities of completing multiple steps, albeit in less than 10% of cells (*48*). In addition to the relatively low efficiency, base editor systems do not have a built-in capability for addressing endogenous genes in the mammalian genome; instead, they require additional synthetic DNAs to be introduced into the cell for a functional gene expression output. By contrast, proGuides displayed high efficiency and syntactic activity for at least seven steps when delivered to mammalian cells as plasmid DNA, and they can be directed to turn on transcription of endogenous genes without requiring insertion of any heterologous DNA or mutation of the host cell genome. As such, the system possesses a We suggest that these new capabilities give proGuide-based cascades unique advantages for the purposes of programming the differentiation of cells through sequential intermediate cell states.

Some characteristics of current proGuide cascades and their use in cells provide opportunities for further development of the system. Considering that the transition between cell states is thought to occur most effectively when it coincides with cell division (*49*), an ideal system might provide new genetic instruction sets (i.e. set of TFs to be expressed) coinciding with cell division. Decreasing the frequency of current proGuide cascades from one step about every 6 hours to every 12 or 24 h would increase the proportion of steps coinciding with the onset of cell division. In addition, it would decrease the number of steps firing during a single cell division and lengthen the duration of the system in cells. Although reducing the efficiency of proGuide conversion with suboptimal CTS sequences reduced the kinetics of cascades, it also reduced the efficiency in terms of numbers of cell completing that step in a cascade, making it undesirable to include in cascade for programming a high percentage of cells. Mechanisms for reduce the speed and increase the duration of the system await investigation, but we suggest that they may involve manipulating the endonucleolytic activity of Cas9 (*15*, *50*, *51*) and using forms of plasmid DNA that are retained in cells (*52*, *53, 54*, *55*).

The advances described in this study have converted proGuides from a proof-of-concept tool to a system that is amenable for programming mammalian cell differentiation. Given the complexity of cellular differentiation and number of intermediate cell types identified by scRNA-seq (*57*), having seven addressable steps for programming cell state changes would be sufficient for deriving most cell types from a pluripotent stem cell starting point. The Cas9-VPR protein has been shown to display transcriptional activation functionality for a wide range of genes, making it possible to activate any gene in the genome with the existing system (*29*). And the use of plasmids to encode the system components provides a substantial advantage for the synthesis and testing of new proGuide cascades in cells. Whereas integrating synthetic DNA elements into a mammalian cell genome can take weeks or months to complete, new proGuide plasmids can be generated from base plasmids in a few days with the ease of simple and inexpensive recombinant DNA techniques that are routinely performed in most molecular biology laboratories. In addition, the generation of new proGuide plasmids is scalable to make hundreds at once, and individual plasmids can be combined to make a variety of mixtures to formulate a large number of distinct cascades encoding the activation of different TFs and different steps. As such, the search space that can be addressed for combinations of TFs for effective differentiation of cells will most likely be limited by the ability to evaluate the effects on cells and not on the production of the proGuide cascade instruction sets. In summary, we suggest that these new proGuides represent new achievements for several parameters, including: syntactic control of instructions, efficiency of progression through multiple steps, low background activity (i.e. leak), programmability to activate any gene in the genome, the number of programmable steps, and an independence from genomic DNA breaks or insertion of heterologous DNA elements into the genome.

### Limitations

Due to the advantages gained from using plasmids to evaluate DNA elements and to build de novo cascades of proGuides, the effects of the current proGuides have been examined exclusively when encoded by plasmid DNA introduced to cells via transient transfection or nucleofection methods. It remains unknown if integrating DNA encoding cascades of proGuides affects functionality; we suggest that integrated cascades will likely have increased duration and slower progression, and display decreased efficiency on a per-step basis because the number of proGuide DNA substrates for each step will be reduced relative to plasmid encoded system. Although empirically measured effects of proGuide variants drove development of current proGuides, effects on the activity of the matureGuide RNA and effects on the conversion from a proGuide were considered separately during development without comprehensive experimental separation of the two effects from each other. As such, if any future optimization of proGuide sequences were to occur, it would be aided by additional experiments elucidating whether improvements are possible for either activity of guide RNA or conversion frequencies.

## Acknowledgments

We thank Matthew MacDougall for initial design of proGuide CTS optimization libraries, Hannah Pennington for development of proGuide CRISPRa evaluations, Andy May for insights into guide RNA engineering, the Rush University Genomics and Microbiome Core Facility (GMCF) team for DNA sequencing, Jose Lopez for expert assistance in generating essential materials necessary for experiments in this study, and Nick Timmins and Jalees Rehman for review and comments on manuscript during the writing process.

## Funding

National Institutes of Health grant R01GM139894 (BJM)

## Author contributions

Conceptualization: AKP, RC, BJM

Methodology: AKP, ACN, CH, MRR, MM, RFD, SA, NGB

Visualization: AKP, ACN, GEM, NGB, BJM Investigation: AKP, ACN, GEM, CH, RFD, SA Funding acquisition: RC, BJM

Supervision: AKP, NGB, RC, BJM Writing – original draft: AKP, BJM

Writing – review & editing: AKP, GEM, NGB, RC, BJM

## Competing interests

RC, NGB and BJM are cofounders of and have equity interests in Syntax Bio; AKP, GEM, ACN, MRR, GEM, RFD, SA, and CH are employees of and/or have equity interests in Syntax Bio. AKP, ACN, NGB, RC and BJM are co-inventors on patent filings related to the proGuide system.

## Data and materials availability

All data are available in the main text or the supplementary materials. Raw DNA sequencing data will be made available upon request. Materials used in the analysis (plasmid DNA) will be available to any not-for-profit researcher for purposes of reproducing or extending the analysis pending completion of material transfer agreement (MTA) with Syntax Bio and the researcher.

## Supplementary Materials

Materials and Methods

Figs. S1 to S15

Tables S1 to S9

## Materials and Methods

### Plasmid DNA cloning

Standard molecular biology methods were used for engineering all plasmids DNA used in this study. A base plasmid for expression of guide RNA was synthesized (GenScript) based on the sequence of the pSpgRNA plasmid (*1*) with modification to enable new spacer sequences to be inserted by Golden Gate cloning with DNA oligos (Genewiz) as previously described (*2*). All new plasmids encoding proGuide RNA variants were synthesized (Genscript) using the same backbone and spacer cloning strategy. Mix & Go competent DH5a (Zymo) were used for transformation of ligation reactions, and plasmid DNA was isolated from 50ml overnight cultures with ZymoPURE II Plasmid Midiprep Kit (Zymo). DNA sequences were verified by nanopore DNA sequencing (Plasmidsaurus).

### Human cell lines and cell culture

Lenti-X 293T (293-LX) cell line derived from Human embryonic kidney 293 cells (HEK293T) were obtained from Takara and used to generate an EGFP-expressing 293-LX cell line. Lenti-X 293T cultures were maintained in high glucose DMEM (Thermo Fisher Scientific) supplemented with 10% FBS (GenClone) and 5 µg/mL of Penicillin/Streptomycin (Thermo Fisher Scientific) and grown in ventilated flasks at 37 °C in a 5% CO2 incubator prior to experiments. Cell cultures were routinely split at a 1:10 ratio every 3 days with dissociation agent TrypLE (Thermo Fisher Scientific).

Human induced pluripotent stem cells (iPSC) derived from primary hepatic fibroblasts obtained from a healthy donor were obtained from American Type Culture Collection (ATCC- HYR0103). Cells were maintained in Essential 8 (Thermo Fisher Scientific) with added supplement kit and 5 µg/mL of Penicillin/Streptomycin (Thermo Fisher Scientific) with a daily change of media. Cells were grown on vessels coated with Vitronectin (Thermo Fisher Scientific) at a working concentration of 0.5 µg/cm^2^ in DPBS (Thermo Fisher Scientific). Once cultures reached 60-70% confluency, they were dissociated with Accutase (Stemcell Technologies), and resuspended in culture media supplemented with 10 uM ROCKi Y-27632 (Tocris) and allowed to grow for 24 h post-passage, after which the media was exchanged for culture media (without ROCKi). Cells were maintained in ventilated flasks at 37 °C in 5% CO2.

### EGFP-293LX cell line production

Lenti-X 293T cultures were maintained in high glucose DMEM (Thermo Fisher Scientific) supplemented with 10% FBS (GenClone) and 5 µg/mL of Penicillin/Streptomycin (Thermo Fisher Scientific). A day before transfection, 1.2x10^7^ cells were seeded into tissue culture-treated 150cm^2^ ventilated flasks in 30mL of culture medium. On transfection day, pseudotyped lentiviral vector was produced by transfection using TransIT-293 transfection reagent (MirusBio) using a third-generation, 4-plasmid system of pMDLg/pRRE plus pRSV-Rev, a transfer vector encoding EGFP driven-off of a spleen focus-forming virus (SFFV) promoter, and the plasmid pMD2-G encoding the VSV-G envelope protein. Each flask received a total of 60 µg of plasmid DNA (30 µg transfer vector, 5 µg VSV-G, 20 µg pMDLg/pRRE, 5 µg pRSV-Rev) and 180 μL of TransIT293. Transfection complexes were allowed to form for 15 min at room temperature in 3mL OptiMEM before addition to the flasks. On Day 2 post-transfection, media in the flasks were exchanged with 30 mL fresh complete media. On Day 3 post-transfection, the crude viral supernatant was collected at 48 h post transfection, and cell debris was removed by centrifugation at 1000 x*g* for 10 min at room temperature. Supernatants were either used immediately or stored at −80°C.

Lenti-X 293T cells were plated and transduced at an MOI of 0.1. Cells were passaged 72 h post-transduction and replated at low density. Cells were collected 7 days post-transduction and sorted by GFP fluorescence on an MA-900 (Sony Biotechnology) to positively select the top 30% of EGFP-expressing Lenti-X 293T cells. Cells were expanded over 2 passages, checked for maintenance of EGFP expression, and cryopreserved in CryoStor10.

### Transfection

For lipofection of Lenti-X 293T, cells were dissociated and plated at a concentration of 3 x 10^5^ cells/mL (for a final volume of 2 mL for a 6-well plate and 300 µl for a 48-well plate) in complete DMEM medium. Plated cells were allowed to grow overnight in the incubator at 37 °C with 5 % CO_2_. DNA mixes of purified plasmid DNA were prepared in Opti-MEM according to the manufacturer’s protocol. TransIT-293 transfection reagent (MirusBio) was added at ratio of 4 µl reagent to 1 µg DNA, incubated at room temperature for 20 min, and finally added dropwise across the well. The transfected cells were maintained at 37 °C with 5 % CO_2_ until collection.

### Nucleofection

Nucleofections of human iPSC were performed using the Amaxa 4D-Nucleofector unit (Lonza, New Jersey, USA). Nucleofector settings had been previously established empirically in the lab (program CB-125, P3 solution) to favor high cell viability and DNA delivery efficiency. Cells were harvested by dissociating cells with Accutase, once detached an equal volume of media (plus 10 µM ROCKi) was added, and cell suspension was mixed. Cell numbers were quantified by NucleoCounter® (Chemometec, Allerod, Denmark). 800,000 cells per nucleovette condition were centrifuged at 300 x*g* for 5 min at room temperature. Subsequently, the supernatant was aspirated and cells were resuspended in P3 solution reagent (10 µL per 800,000 cells). DNA plasmids for nucleofection mixes were prepared in MilliQ water. Individual nucleovette wells of a 96-well plate condition received a total of 1µg DNA mass at 250 ng/µL concentration and 6 µL of P3. A 10 µL aliquot of the P3 cell suspension was added to each nucleovette prior to nucleofection. After executing the nucleofection program, cells were incubated at room temperature for 10 min and then resuspended with 80 µL of media (with 10 µM ROCKi) for a total volume of 100 uL per nucleovette. Cells were seeded at 120,000 cells for 96-well and 170,000 cells for 48-well formats. To decrease apoptosis in transfected iPSC, 100 µM Z-VAD FMK was added to the media for the first 24 h, and a BCL-XL expression plasmid was included in the DNA mix (*3*). Post-transfection, cells were incubated for 24 h at 37 °C with 5% CO_2_ in a humidified incubator.

### Flow Cytometry

Single-cell suspensions were prepared by detaching the cells with TrypLE and resuspended in DMEM media. Subsequently, cells were washed in the following cell staining buffer (CSB): 2% FBS, 0.5 mM EDTA, and 0.4% NaN_3_. For CXCR4 and CD105 detection, cells were stained with Brilliant Violet 421 anti-human CD184 (CXCR4) and APC anti-human CD105 at a 1:100 concentration. For DLL4, CD4, CD105 detection, cells were stained with PE/Dazzle 594 anti-human CD4 (1:600), PE anti-human DLL4 (1:200), and Brilliant Violet 421 anti-human CD105 (1:600), all from BioLegend (San Diego, CA, USA). Antibody incubation was for 30 min at 4 °C in the dark. Flow cytometry antibody staining for iPSCs included additional steps: After iPSC were dissociated and washed once, they were incubated for 20 min on ice with an Fc-blocking step using IgG human serum diluted 1:400 in CSB. The iPSC were centrifuged at 500 xg for 5 min, followed by decantation, and addition of antibody solution. For the live-dead stain, cells were resuspended in CSB containing DRAQ7 (1:1000) or with Propidium Iodide at 1 µg/mL. Cells were analyzed with either a Cytoflex (Beckman Coulter) or NovoCyte Penteon (Agilent) flow cytometer. Data analysis was performed using FlowJo v10.10.0. At least 3x10^5^ singlet, live cells were counted for each sample.

### Generation of virtual library of non-genome targeting CTS

See **Fig. S11** for schematic illustration of the overall process. Ensembl versions of genomes at release 103 were acquired for human (homo_sapiens), mouse (mus_musculus), cow (bos_taurus), atlantic salmon (salmo_salar), chicken (gallus_gallus), pig (sus_scrofa). The pacific bluefin tuna genome was acquired from the NCBI with Genbank accession number GCA_009176245.1 (thunnus orientalis, tuna2). Blast databases were created for each genome using ncbi-blast version 2.12. A fasta of two million random CTS sequences was created using RandomSeq (*4*) with settings FLS, n=2000000, cap_start=False, cap_stop=True, fasta=True, allow_start=False, allow_stop=True, stop_codons=’AGG,TGG,CGG,GGG’, length=20. This generates a compendium of 23 bp CTS with the 4 potential NGG Cas9 recognition sites at their 3’ ends. NCBI blastn was run against these genomes sequentially with the 2M CTS using default settings and task ‘blastn-short’ which is optimized for sequences shorter than 50 bases. Using data from the foundational GuideSeq paper (*5*) as a reference point bona-fide off target sites “*harbored as many as six mismatches within the protospacer sequence*”. Hence when CTS sequences were aligned to each genome they were only included in the next alignment if they had 7 or greater mismatches and additionally less then a 16 base stretch of match. Additionally, a set of heuristics based on literature study of off-target analyses were used to filter these sequences further. Sequences with higher G-content in the non-seed region distal from the PAM motif have been shown to have stronger off target activity (*6*). Hence, all sequences with greater than or equal to 5 G’s in the 10bp most distal to the PAM were discarded. Thymines in the 3’ region of spacer are disfavored for both Cas9 loading and cleavage efficiency(*7*), hence all sequences with 3 or more T’s in the region directly adjacent to the PAM were discarded. gRNA with extremes of GC content “tend to be nonfunctional”(*7*), using this reference and the GC content of the improved gRNA screen in (*8*) as a guide only those CTS with GC content of between 35 and 75% were kept. Additionally, our backbone plasmid contains 1EcoRI, 2 BsmBI, and 1 SapI that are used for our cloning, any CTS that generated additional instances of these RE sites within the backbone were discarded. At the conclusion of alignment and filtering we were left with a of list of >50,000 CTS sequences.

### RNA Sequencing of guide RNAs

GFP-293LX cells in 6-well plates were transfected with Cas9 and a proGuide, with or without the trigger guide. Thirty-six hours post-transfection, cells were detached and counted. One million cells were directly lysed in 300 µL of Tri-Reagent (Zymo Research) and stored at -80 °C. Total RNA was extracted using the Direct-zol™ RNA Miniprep Plus kit (Zymo Research) according to the manufacturer’s guidelines. cDNA of the guide RNAs was synthesized with the Template Switching RT Enzyme Mix (NEB) using the 2nd Strand cDNA Synthesis Protocol. Specificity of guide RNA synthesis was ensured by using a 3’ primer specific for the 3’-end of the guide RNA scaffold present in all fully transcribed guide RNAs. Briefly, 800 ng of total RNA was used for annealing of the 3’-primer in a total volume of 24 µL and incubated for 5 minutes at 70 °C. For reverse transcription, 9 µL of the annealing mixture was added to 6 µL of the RT mix and incubated at 42 °C for 90 min, followed by 5 min at 85 °C. For the subsequent synthesis of the 2nd strand, 10 µL of the RT mix was added to 90 µL of the 2nd strand cDNA synthesis reaction mix and incubated at 37 °C for 15 min, then at 95 °C for 1 min, and 65 °C for 10 min. Finally, 30 µL of this mixture was cycled for 20 PCR cycles to amplify the cDNA for the guide RNA, and sequenced using an Illumina MiniSeq, generating 150 bp paired-end reads.

Fastqs of RNAseq data following PCR of guide sequences were assembled and R1 and R2 filtered to remove triggerGuide sequences (which share the same scaffold sequence). Reads were also filtered to remove both for very short (<50 bp) and very long reads (>160 bp). Reads were modified with cutadapt to remove both CS1 (ACACTGACGACATGGTTCCTACAGGG) and CS2 (AGACCAAGTCTCTCGTACCGTA) linkers from the reads allowing 1 mismatch. Flash was used to combine R1 and R2 with min-overlap 10. Crispresso2 was run on all architectures (See Table S2 for settings). Perfect adjacent repair is a heuristic of repair outcomes that are “close” perfect repair but owing to sequencing error or slight differences in size. A perfect adjacent repair has at most 5 substitutions, 1 and only one deletion within 3bp of a perfect repair, and up to 4 additional deletions of which all must be under 5bp, and only 2 can be above 2bp. Inversions were detected with crispresso2 using the architecture if the excised DNA was placed in reverse in a perfect repair.

---

**Fig. S1.**
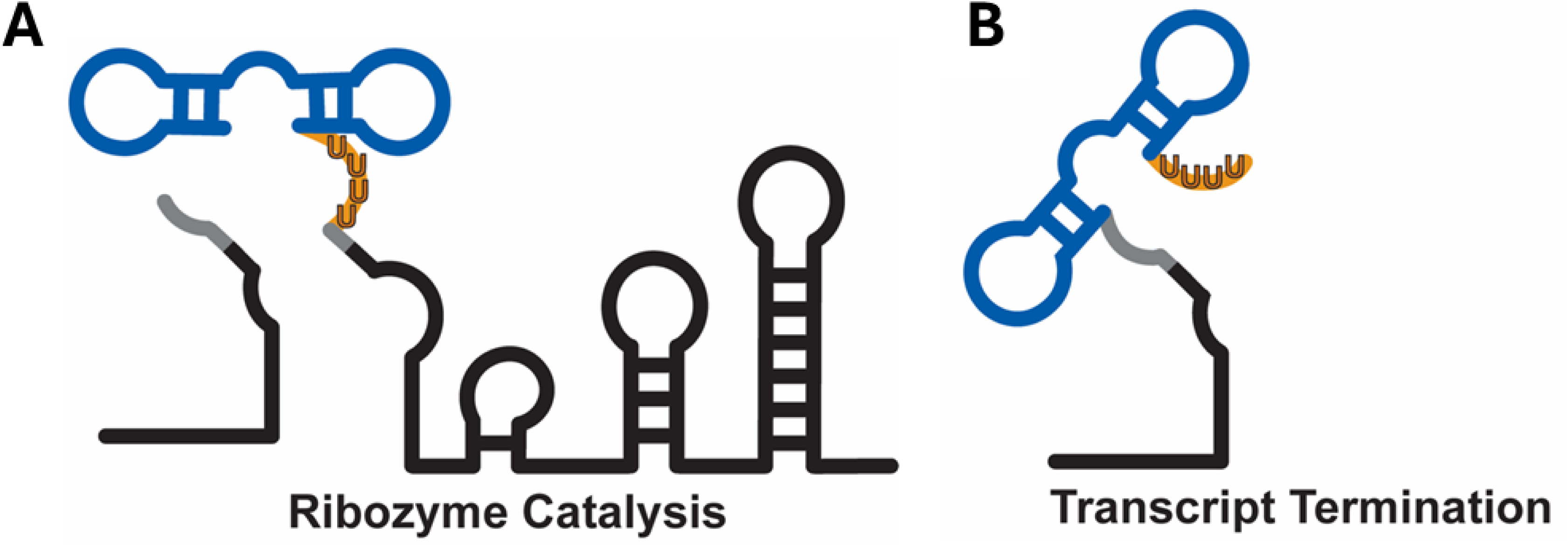
Schematic of previous combined methods for inactivating RNA to make proGuides. Illustrations of the RNA transcripts resulting from the redundant inactivation elements incorporated into a tetraloop region (gray). Inactivation elements encoded a hammerhead ribozyme (blue) and an RNA Pol III termination signal of six contiguous thymidine residues (orange). **A**) Ribozyme-based inactivation method utilizes cleavage of the hammerhead ribozyme such that the upstream and downstream sequences dissociate into two separate RNA strands. **B**) The transcript termination method prevents transcription of necessary functional elements of the guide RNA (i.e. hairpin 1, hairpin 2) by placing a termination signal upstream of those elements in the proGuide.

**Fig. S2.**
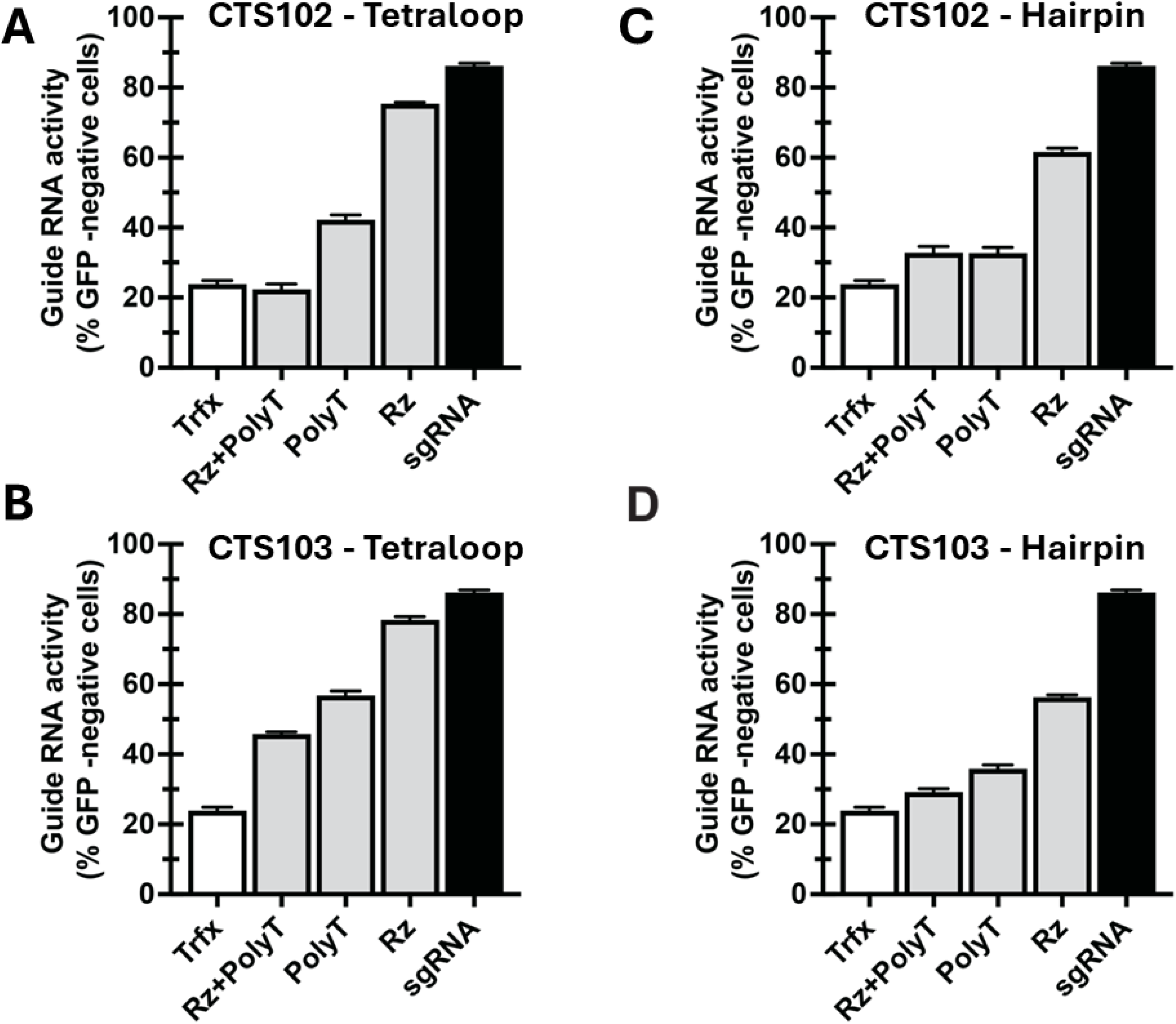
Poly T tract alone is more effective than ribozyme alone at inactivating proGuides. Data are similar to those described in Fig. 1C with the exception that different CTS flank the inactivation moiety. **A,B**) CTS102 and CTS103 flank the inactivation moiety within the gRNA tetraloop, respectively. **B,C**) CTS102 and CTS103 flank the inactivation moiety within hairpin 1 of the gRNA, respectively. Residual activity of proGuides in cells was primarily affected by the inactivation unit (ribozyme vs. polyT tract) and insertion location (hairpin vs. tetraloop) and minimally affected by the CTS sequence.

**Fig. S3.**
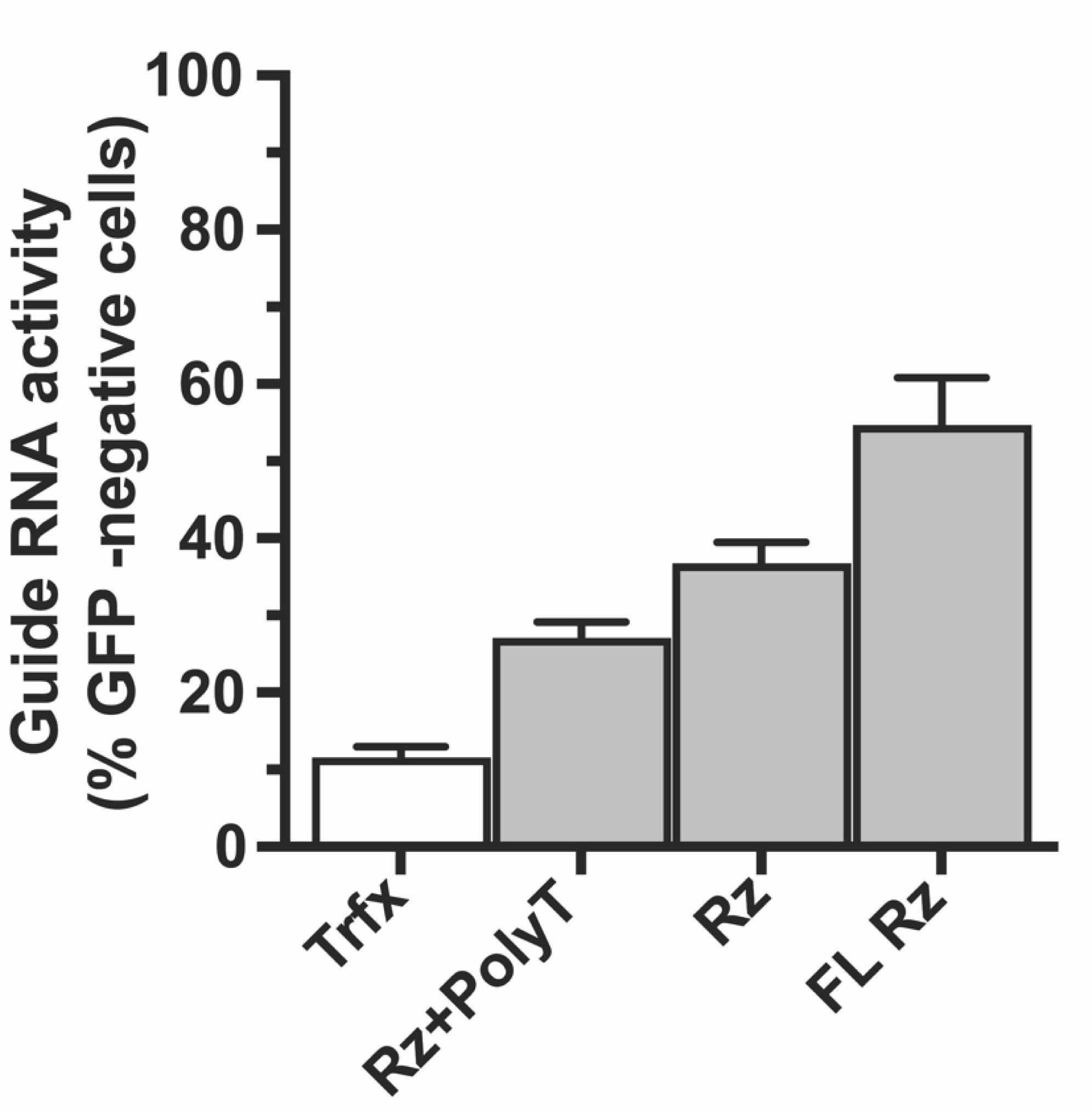
A full-length hammerhead ribozyme is ineffective as an inactivating unit for proGuides. Similar to experiments shown in Fig 1C. For a given proGuide, different inactivation units generate different levels of leak of GFP disruption in the absence of a trigger guide. Ribozyme only inactivation units led to a larger level of leak. The full-length hammerhead Rz, which exhibits higher *in vivo* nucleolytic activity (REF), displayed the highest level of unwanted proGuide activity.

**Fig. S4.**
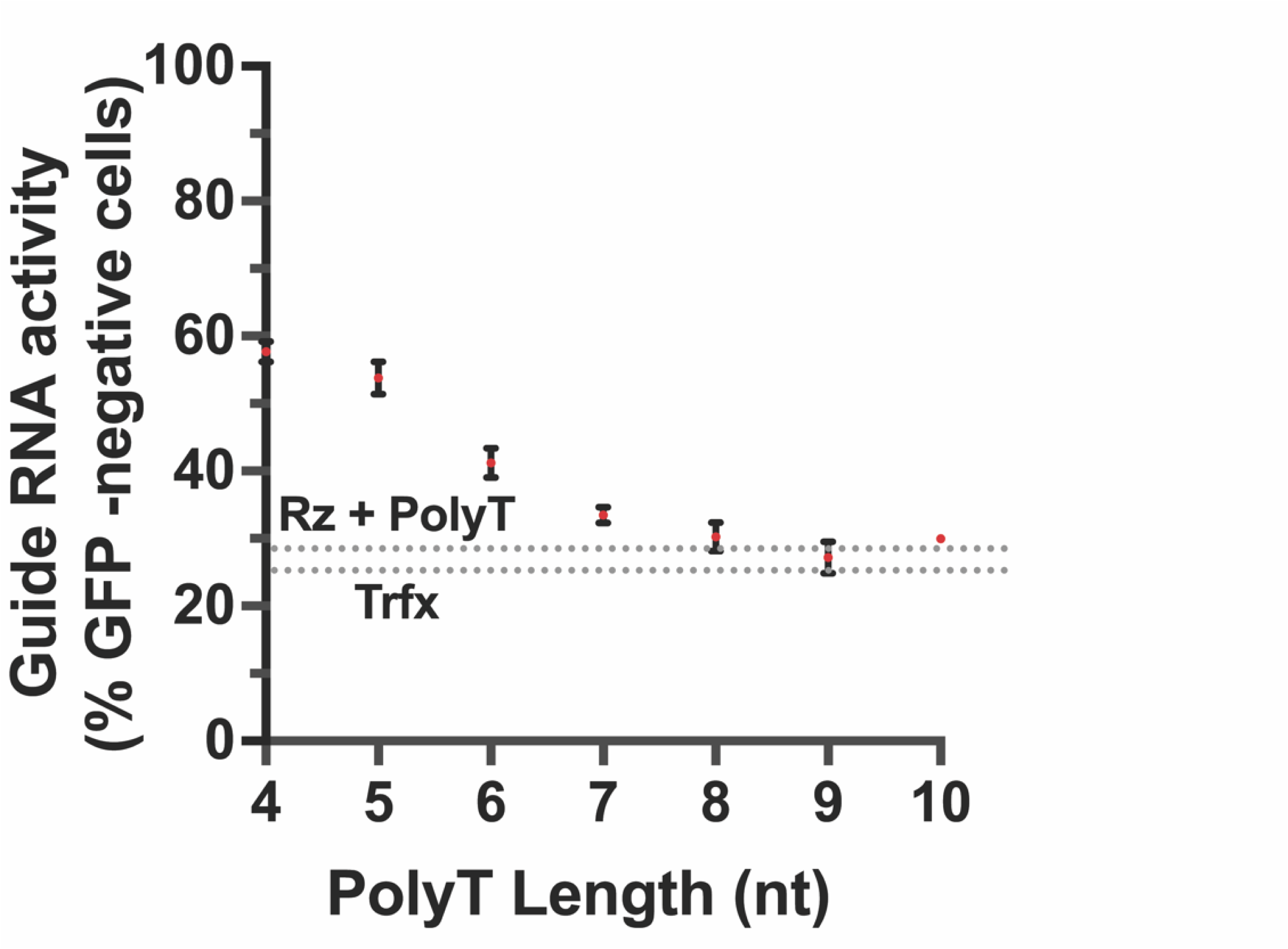
Increasing length of the polyT tracts embedded in hairpin 1 reduces proGuide leakiness. Similar to experiment shown in Fig 1D, except polyT tracts were inserted into hairpin 1 instead of into the tetraloop.

**Fig. S5.**
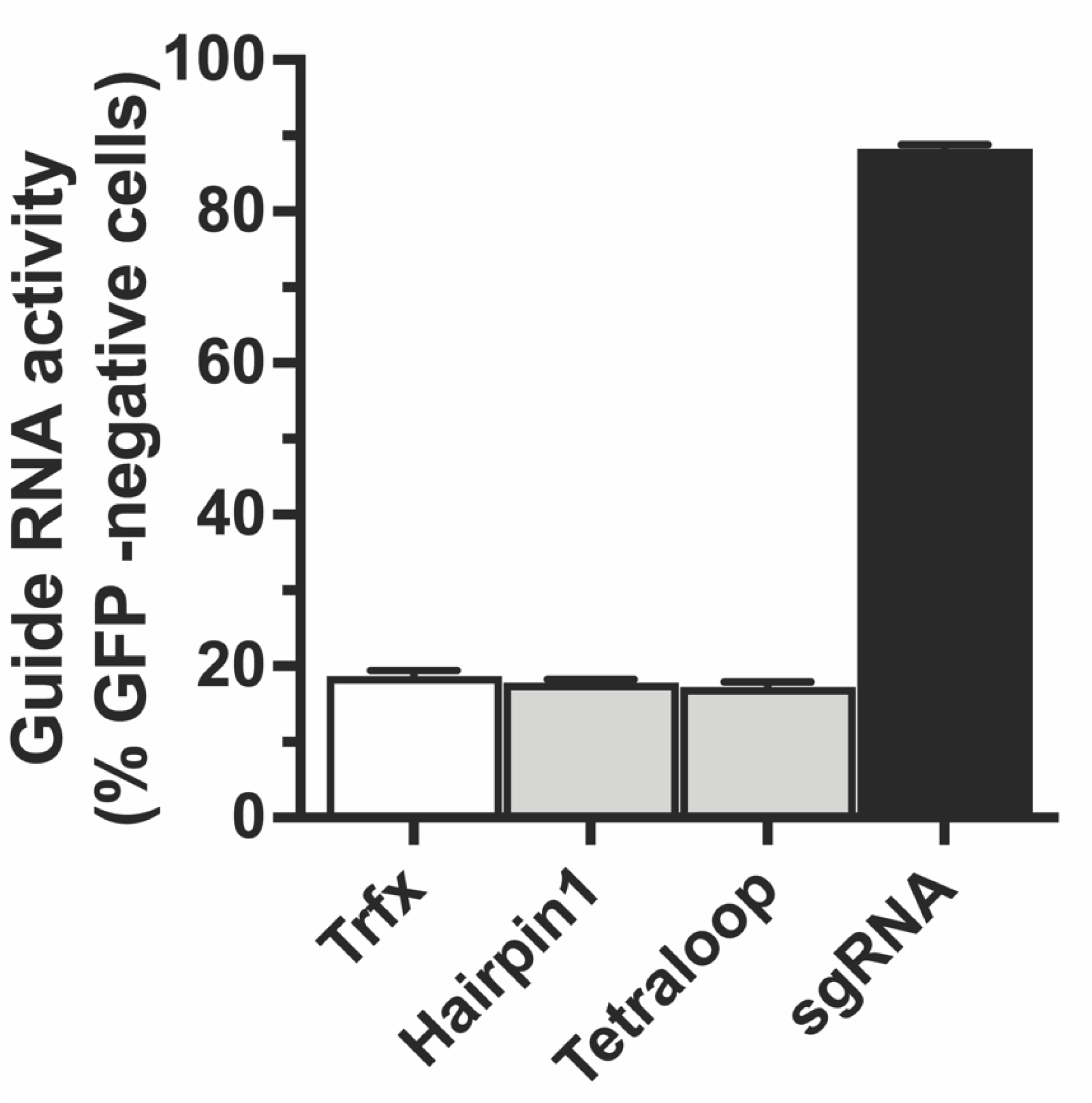
Early terminated guide RNAs are inactive. RNA transcripts comprising only the 5’ portion of the proGuide sequence upstream of the polyT termination tract at either the tetraloop site or the hairpin 1 site (gray bars) exhibit no residual Cas9 cleavage-competency in cells compared to transfected cells (white bar).

**Fig. S6.**
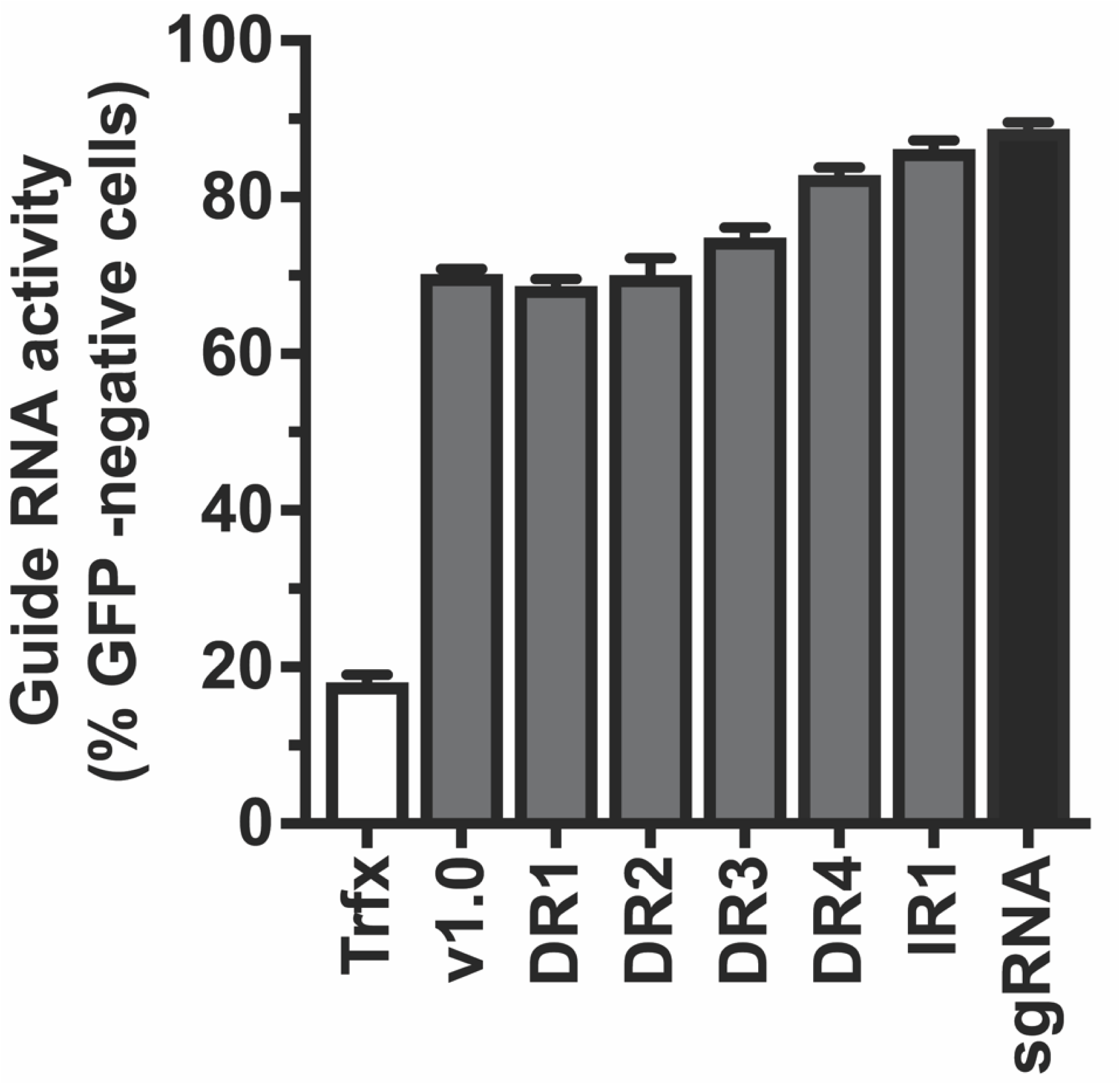
The inverted repeat CTS configuration confers efficacy similar to an sgRNA. Similar to experiment shown in Fig. 2B, except inactivation sequences were inserted into the hairpin 1 site instead of the tetraloop of proGuide constructs.

**Fig. S7.**
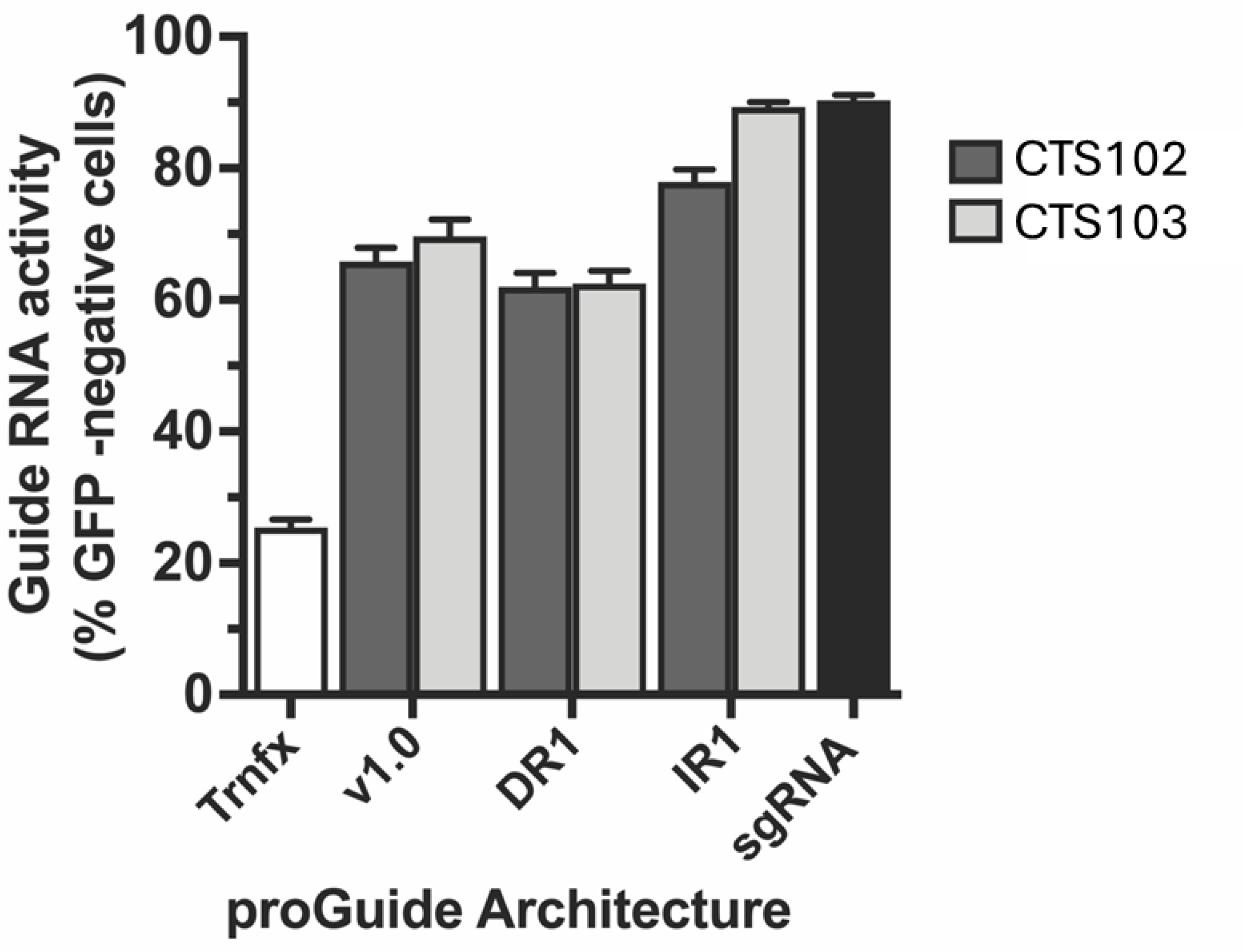
Inverted repeat configuration outperforms direct repeat configuration for multiple CTS sequences. Similar to experiment shown in Fig. 2B, except proGuides had CTS102 and CTS103 sequences instead of CTS101.

**Fig. S8.**
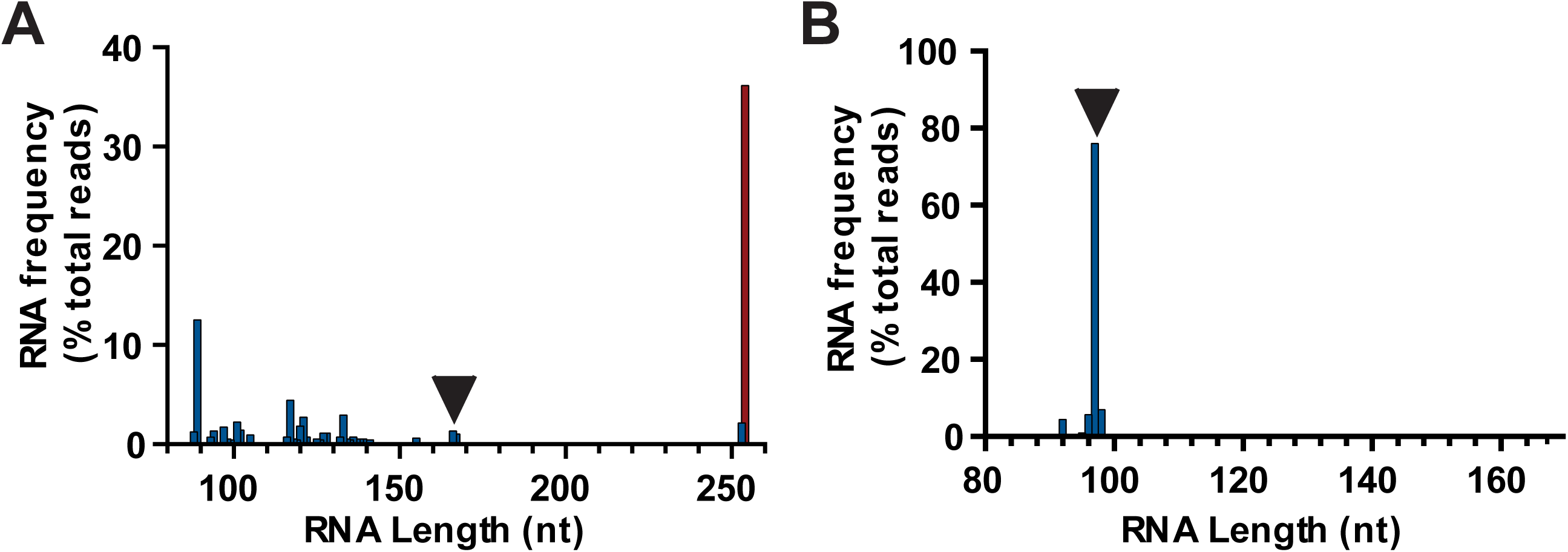
Analysis of RNA produced from proGuide plasmids following co-transfection with trigger sgRNA. **A)** RNA was isolated from HEK293T cells transfected with a trigger sgRNA, Cas9 plasmids and the previously published proGuide architecture containing a ribozyme and CTS102 sites. Oligos specific to the proGuide spacer sequence and the 3’ end of the common guide RNA sequence were used to amplify cDNA made from isolated RNA. DNA sequencing of amplicon products were analyzed with Crispresso2, and the graph shows the distribution of fragment lengths that mapped to the proGuide sequence. The arrowhead indicates the size of an expected perfect repair outcome, where the ends of the two CTS are repaired via NHEJ. **B)** Same procedure as in (A), except the proGuide harbored the polyT tract inactivation element and higher efficiency CTS101 sites in an inverted repeat orientation.

**Fig. S9.**
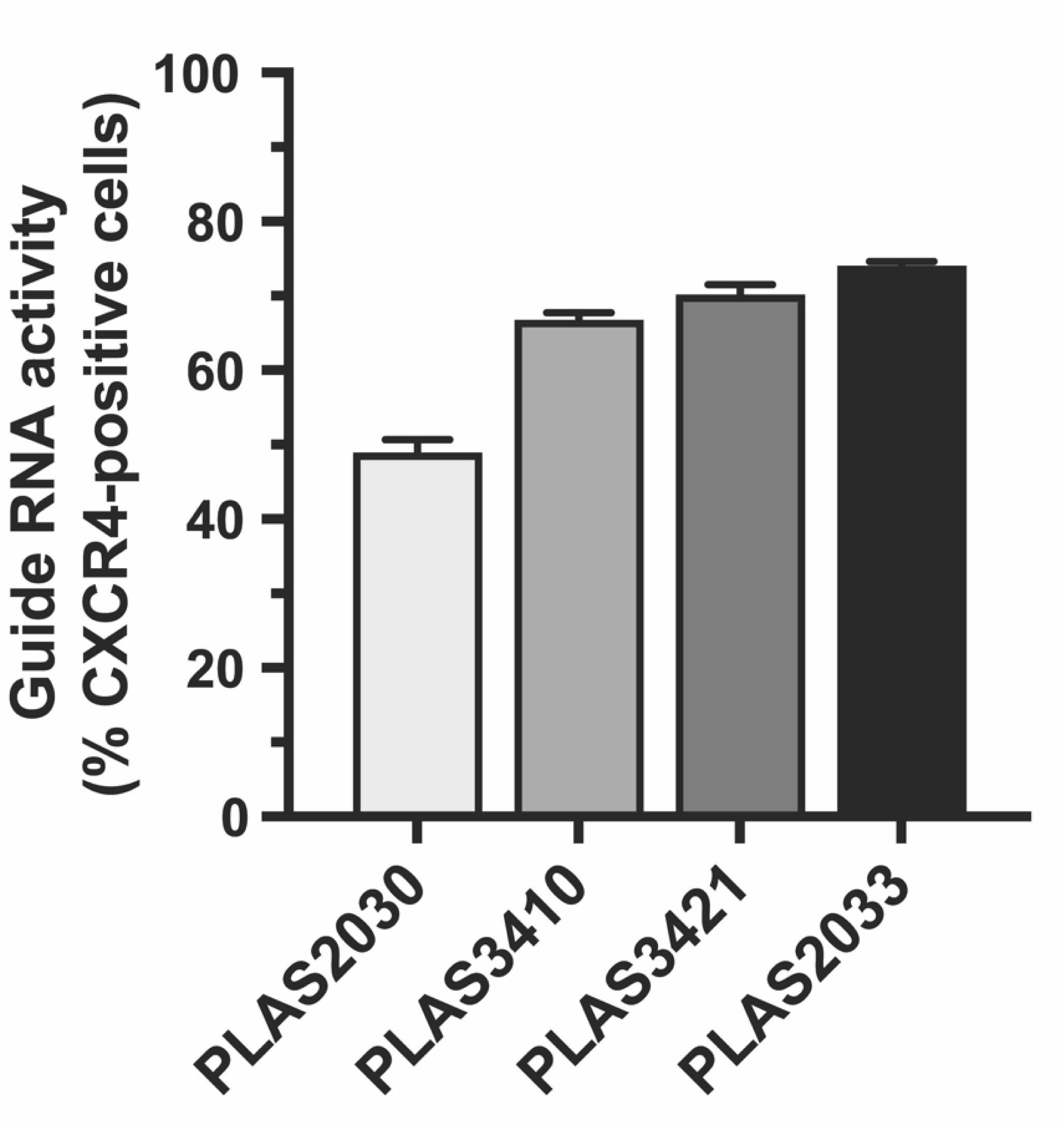
Evaluation of CXCR4-targeting 14 nt spacer sequences for activation of CXCR4. Frequency of cells expressing CXCR4 protein on the cell surface was measured by flow cytometry 48 hr after transfection with plasmids encoding Cas9-VPR and a CXCR4-targeting sgRNA. Each guide RNA plasmid used a different 14 nt sequence for the spacer targeting CXCR4 promoter region. PLAS2033(GGAAGGAGGGCGGCA), PLAS3410(CTGCTGTTTGCGGG), PLAS3421(AACGCGTCTCTCTG), PLAS2030(GCGGGGAATGGCGT)

**Fig. S10.**
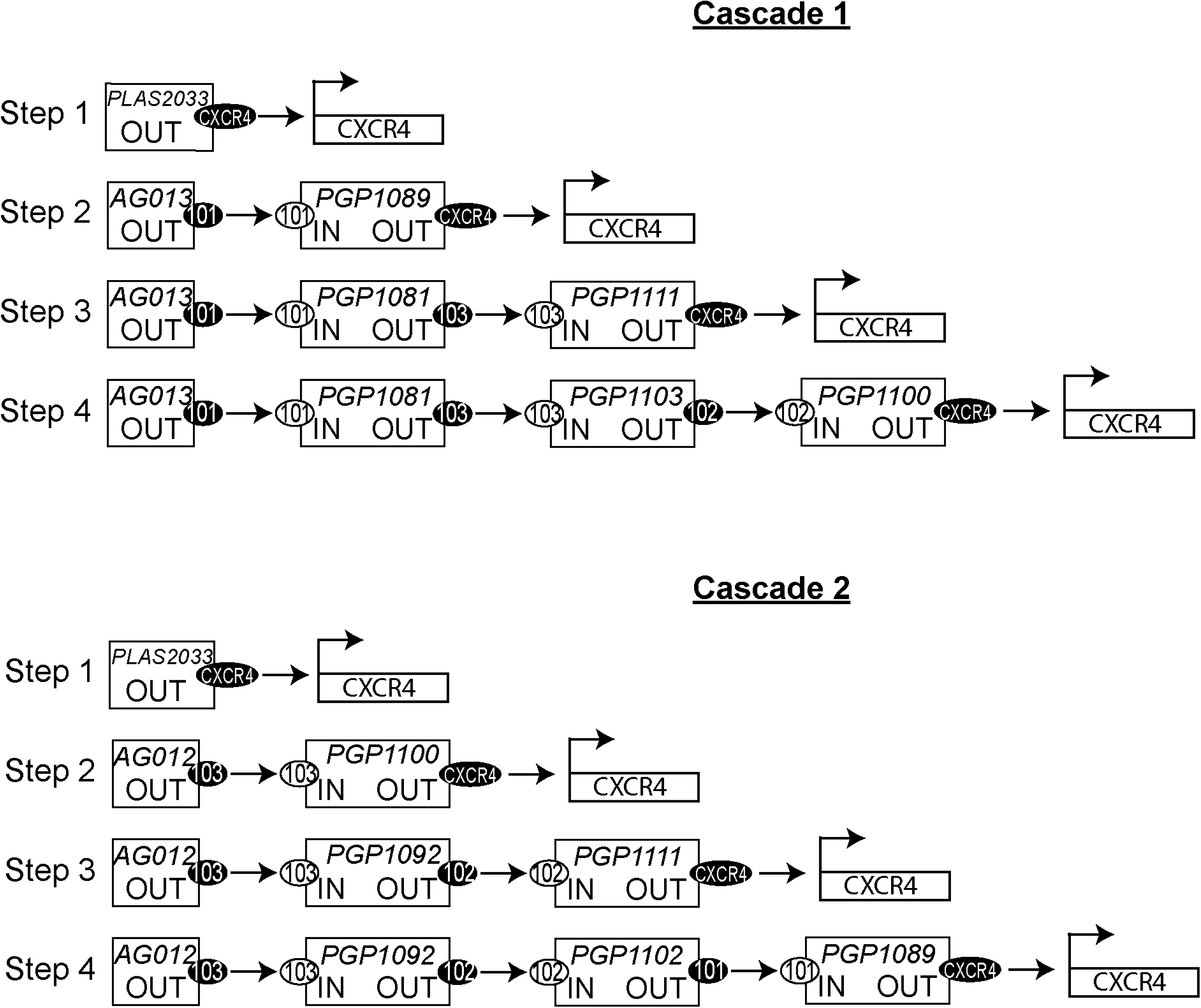
Schematic of proGuide cascades resulting in the transcriptional activation of an endogenous gene. Following the design of the cascade schematic illustrated in Fig. 1A, each row depicts the arrangements of plasmid DNA delivered via transient transfection to HEK293 cells for graphs shown in Fig 3C. PLASxxxx and PGPxxxx rubric corresponds to plasmid DNA unique identification labels for guide RNA and proGuide expression plasmids, respectively. The numbers within ovals (101, 102, 103) refer to the CTS sequences (white) and spacer sequences (black) present in each plasmid DNA. The black oval containing CXCR4 indicates the presence of a 14nt spacer sequence targeting the CXCR4 promoter region for CRISPRa transcriptional activation by Cas9-VPR.

**Fig. S11.**
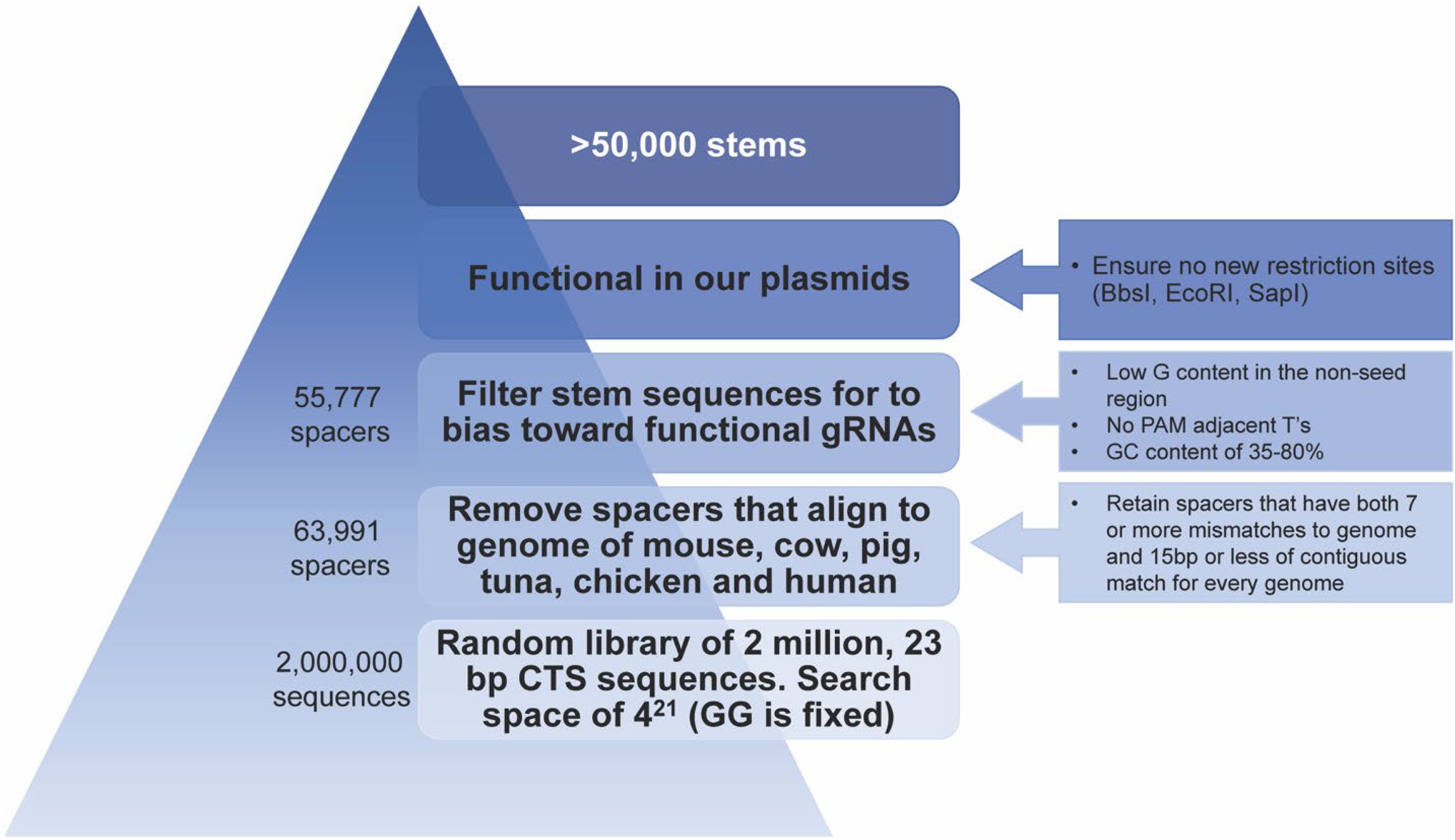
Schematic for the process of generating virtual library of CTS sequences for proGuides. See Materials and Methods for description.

**Fig. S12.**
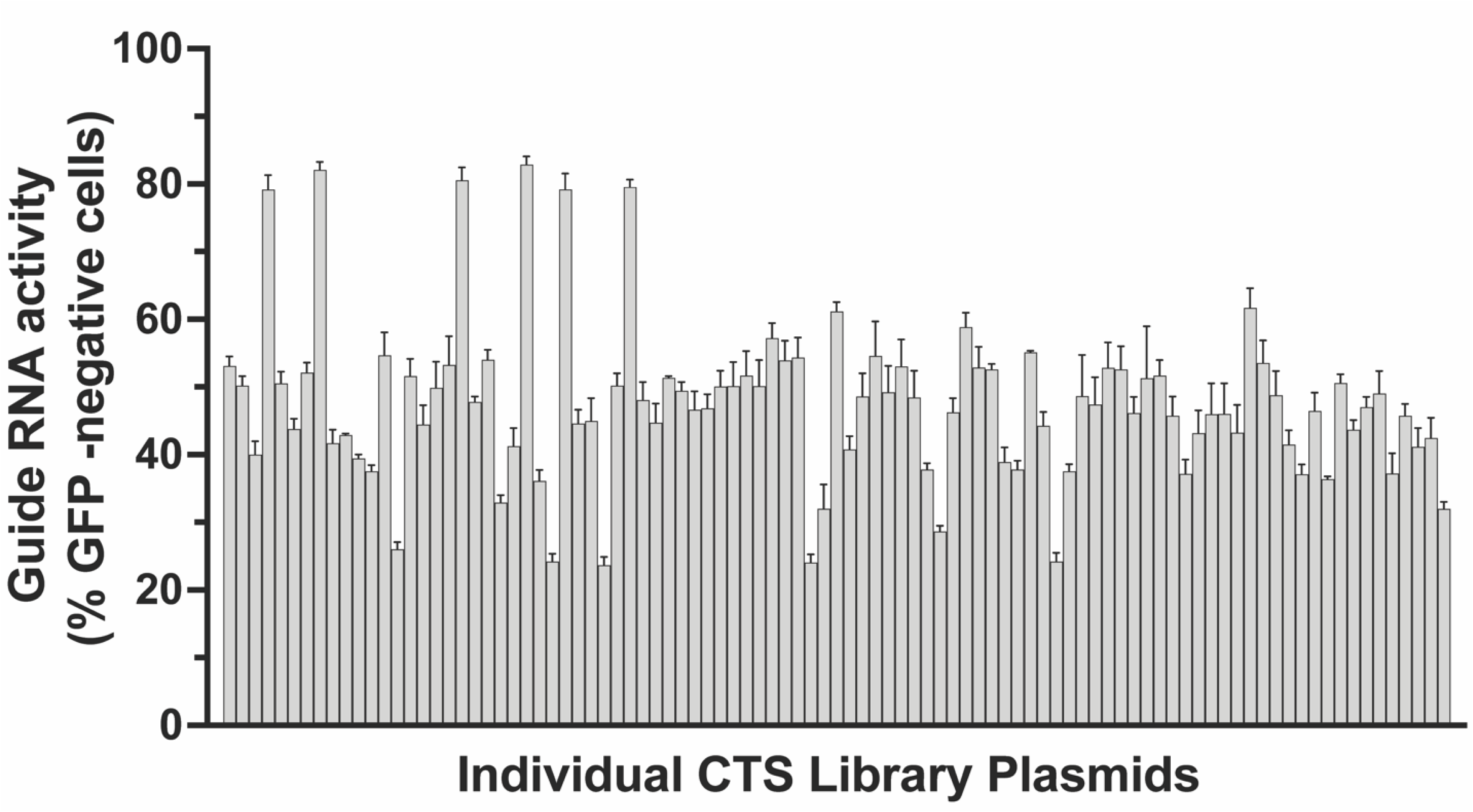
Screen of 96 CTS sequences for conversion to an active guide RNA. Individual plasmid DNA expressing both an EGFP-targeting proGuide and a trigger guide RNA targeting the CTS sequences in the corresponding proGuide were constructed. A plasmid library of approximately 10,000 sequences from Fig. S10 was generated, from which 96 individual plasmid DNA were isolated and evaluated. Disruption of EGFP was measured two days after transient transfection. Note that DNA sequencing of plasmids showed that the six very high activity plasmids were defective proGuide cloning artifacts, and the five very low activity plasmids harbored defective trigger RNAs. Excluding these sequences, the CTS sequences exhibiting the highest GFP knockdown were subsequently used to construct proGuides evaluated in Fig. 3D.

**Fig. S13.**
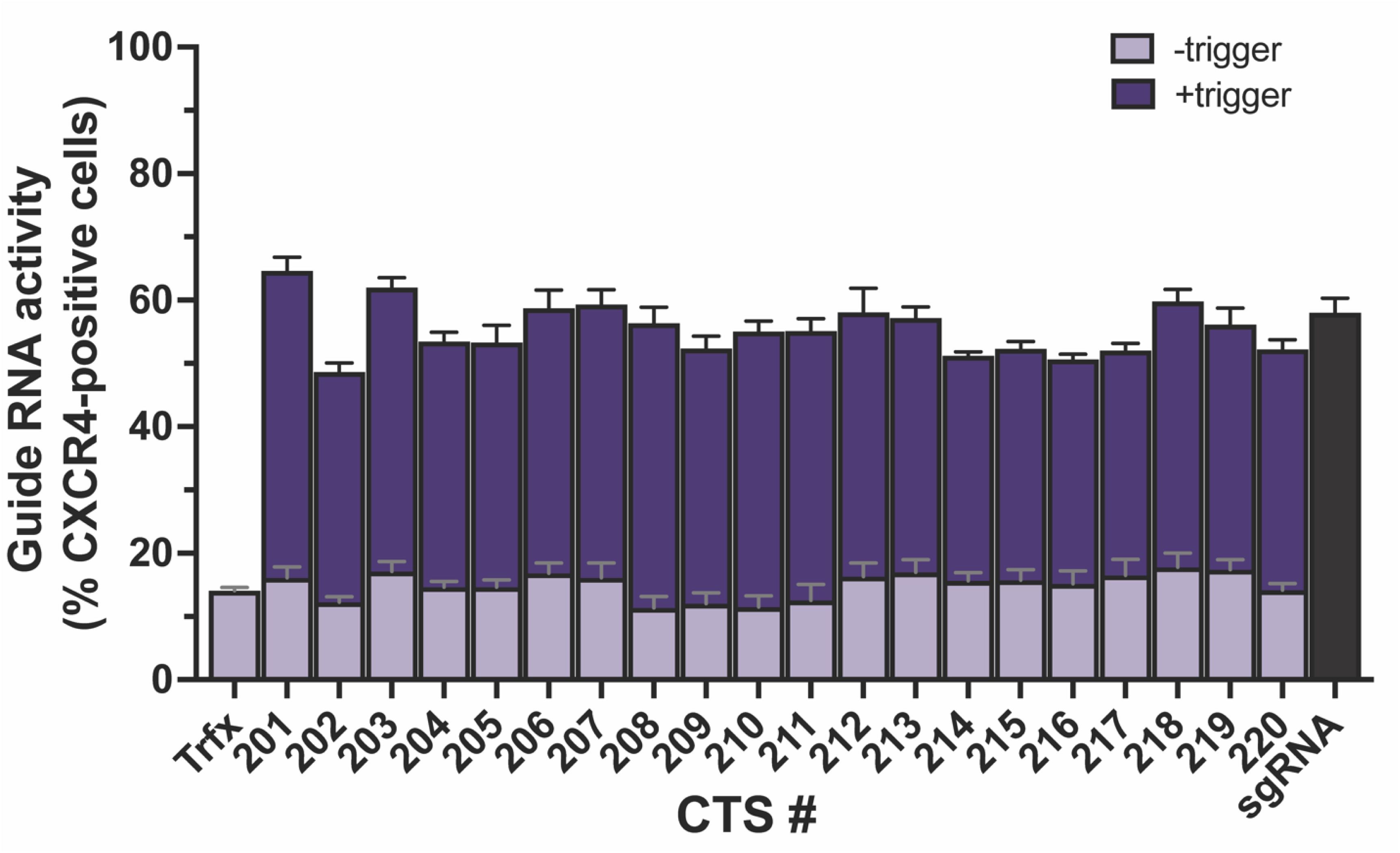
Evaluation of top CTS for proGuides using CRISPRa activation of CXCR4 surface protein expression. Related to Fig. 3D, except proGuides contained 14 nt spacer sequences targeting the CXCR4 promoter. Transient transfection of the proGuide plasmids and a Cas9-VPR expression plasmid enabled increased CXCR4 expression only with expression of the trigger gRNA.

**Fig. S14.**
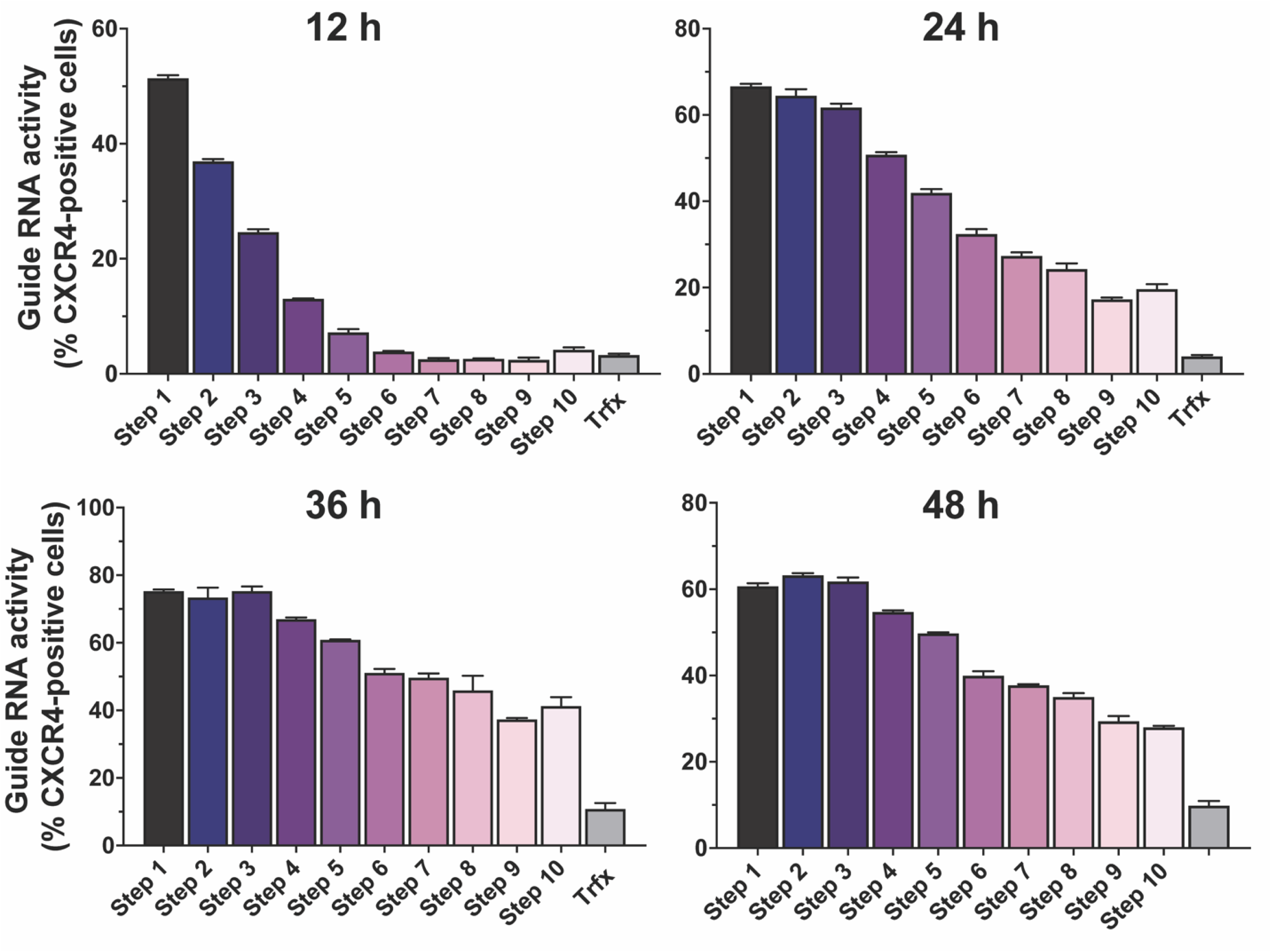
Evaluation of kinetics of a 10-step cascade of proGuides, The forward cascade from **Fig 3E** was transfected into HEK293T cells with a Cas9-VPR and a proGuide for activation of CXCR4 at the indicated step. Flow cytometry for surface CXCR4 protein expression was performed at the indicated times after transfection.

**Fig. S15.**
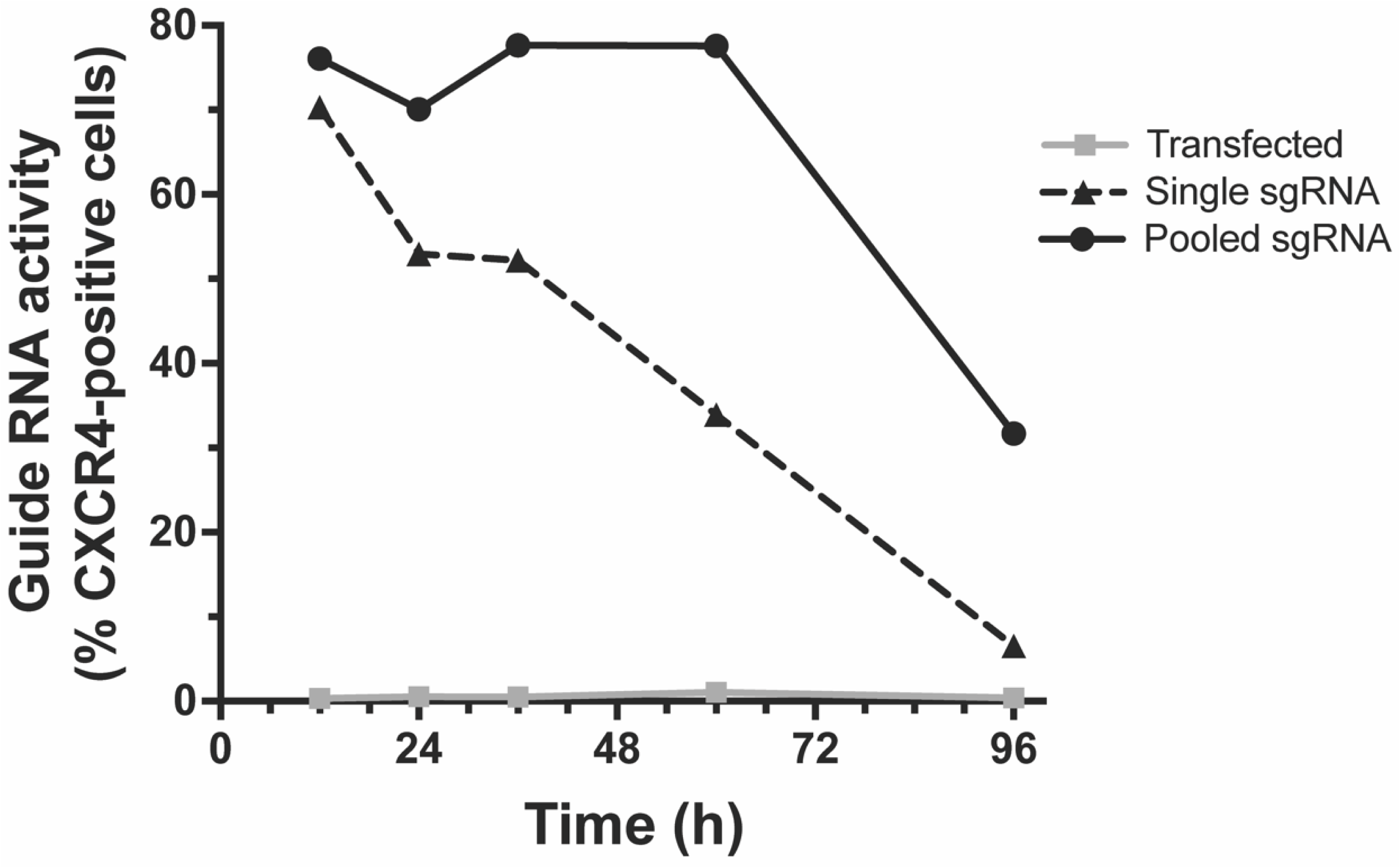
Pools of guide RNA generate increased and sustained target activation. A pool of four sgRNAs (**Fig. S9**) that target mulitple regions of the CXCR4 promoter had higher and more sustained activation than from a single sgRNA even when the total DNA mass was kept constant.

